# Interferometric imaging of thermal expansion for temperature control in retinal laser therapy

**DOI:** 10.1101/2021.11.23.469540

**Authors:** David Veysset, Tong Ling, Yueming Zhuo, Daniel Palanker

**Affiliations:** Hansen Experimental Physics Laboratory, Stanford University, Stanford, CA 94305, USA; Department of Ophthalmology, Stanford University, Stanford, CA 94305, USA; Department of Electrical Engineering, Stanford University, Stanford, CA 94305, USA

## Abstract

Precise control of the temperature rise is a prerequisite for proper photothermal therapy. In retinal laser therapy, the heat deposition is primarily governed by the melanin concentration, which can significantly vary across the retina and from patient to patient. In this work, we present a method for determining the optical and thermal properties of layered materials, directly applicable to the retina, using low-energy laser heating and phase-resolved optical coherence tomography (pOCT). The method is demonstrated on a polymer-based tissue phantom heated with a laser pulse focused onto an absorbing layer buried below the phantom’s surface. Using a line-scan spectral-domain pOCT, optical path length changes induced by the thermal expansion were extracted from sequential B-scans. The material properties were then determined by matching the optical path length changes to a thermo-mechanical model developed for fast computation. This method determined the absorption coefficient with a precision of 2.5% and the temperature rise with a precision of about 0.2°C from a single laser exposure, while the peak did not exceed 8°C during 1 ms pulse, which is well within the tissue safety range and significantly more precise than other methods.

## 1. Introduction

Controlling the temperature rise in tissue during laser therapy is fundamental to many medical procedures, from thermal ablation of tumors to dermatological and retinal treatments. Retinal photocoagulation, for instance, has been a standard procedure for decades to treat a number of retinal pathologies, such as proliferative diabetic retinopathy [1], macular edema [2], central serous chorioretinopathy [3], and others [4]. Extent of heating and the induced therapeutic effect depend on the temperature course (magnitude and duration), which, in turn, depends on the laser parameters (wavelength, power, duration, spot size, etc.) and the tissue properties (optical absorption and scattering, heat capacity and conductivity). Recent studies have suggested that the therapeutic window, typically quantified by the Arrhenius integral Ω, of the non-damaging hyperthermia is narrower (0.1 < Ω < 1) than in conventional photocoagulation [5–7], and thus such treatments require higher precision in the temperature control.

In the retina, light absorption is primarily governed by the pigmentation in the retinal pigment epithelium (RPE) and choroid, associated with an optical absorption that is proportional to the melanin concentration [8]. Measuring the optical absorption in the retina (and in most tissues) *in vivo* is, however, particularly challenging since there is no direct optical access to the transmitted light, and it must therefore be done in reflection. Furthermore, the melanin concentration can vary locally within the retina and from patient to patient by a factor of two to four [9]. These variations, if not accounted for, can result in significant temperature differences and thus undesired lesions or sub-therapeutic exposures. To ensure the laser exposure be within the non-damaging therapeutic range (0.1 < Ω < 1), one can estimate using previously published finite-element thermal models of the retina [8,10] that the absorption coefficient must be determined with accuracy no worse than ±20% (see the Non-damaging treatment range section in the Supplemental Document).

To assess the retinal absorption in reflection geometry, one can measure the tissue response to heating. A few approaches have been developed along these lines. One of them, an optoacoustic method, is based on the detection of the pressure waves at the cornea, generated by the expansion of melanosomes during repeated nanosecond laser exposures [11–14]. During the thermal and acoustic confinement, the detected pressure is proportional to the absorbed energy density, related to the absorption coefficient, and the Grüneisen parameter. Precision of the temperature measurement in this approach has been limited to a few degrees. Further, the signal is sensitive to the position of the piezo sensor on the cornea and hence the measurements require a careful calibration to convert the measured pressure waves into temperature. Another approach is based on detecting the changes in back-scattered light due to microbubbles formed by vaporization, which lead to the RPE cell death [15]. However, due to associated tissue damage, this technique is not suitable for non-damaging therapies. Optical coherence tomography (OCT) has also being explored as a non-contact, high spatial (3D) resolution imaging method for *in-situ* monitoring of the tissue deformation [10,16–19]. In particular, phase-sensitive OCT allows tracking of the nanometer-scale optical path length changes, and has been proposed for precise temperature control during photothermal therapy [16]. Yet, to date, experimental investigations have not directly and quantitatively related the phase signal to thermo-mechanical changes in tissues, from which the temperature could be derived. Instead, temperature profiles have been separately estimated via the optoacoustic method mentioned above [16] or analytically, using an *a priori* value for the absorption coefficient [17,18]. The calculated temperature profiles were then qualitatively compared with the phase profiles recorded via OCT, without any direct validation. Recent advances in pOCT demonstrated capabilities of detecting nanometer-scale deformations with sub-ms temporal resolution [20,21], which should enable rapid and precise optical thermometry.

In this study, we present a methodology to precisely determine the optical (absorption coefficient) and thermal (heat conductivity and coefficient of thermal expansion) properties of layered materials using line-field phase-resolved optical coherence tomography (pOCT). In this approach, we monitored the vertical displacements and the changes in the optical path lengths (ΔOPLs) of reflective and scattering layers with nanometer precision in a non-damaging regime. We developed an analytical model to the coupled thermo-elastic problem and solved it in the Hankel-Laplace transformed domain. This approach significantly increases the computation speed compared to finite element simulations, and therefore allows faster fitting of the material properties to match the spatial and temporal aspects of the experimental ΔOPLs.

## 2. Materials and methods

### 2.1. Optical setup

The optical layout (Fig. 1) followed the design by Pandiyan et al. [22] for high-speed line-scan OCT imaging. We here summarize the main components of the design; further details on the optical system can be found in Ref. [22]. The illumination is provided by a fiber-coupled super-luminescent diode (M-T-850-HP-I, Superlum, 9-mW power) with a center wavelength of 840 nm and a bandwidth of 130 nm, providing an axial resolution of 2.4 μm in air. After collimation with a reflective collimator (RC) to a diameter of 4 mm, the beam is shaped using a cylindrical lens (CL1) to yield a line-illumination at the tissue phantom, which is held around its periphery and whose front and back surfaces are in contact with air. After CL1, the beam is split between the reference and sample arms with a 30:70 (R:T) beam splitter. In the sample arm, the beam, after going through magnification telescopes, is focused to a line field of 0.8-mm in length (FWHM of the Gaussian intensity profile). A galvo-scanner is used to steer the beam laterally (*y* direction) at the sample surface and a deformable mirror is used for fine focus adjustment and for the future adaptive optics implementation. The entrance pupil plane (P), the galvo-scanner, and the deformable mirror are optically conjugated using afocal telescopes (L1+L2, L3+L4).

**Fig. 1.**
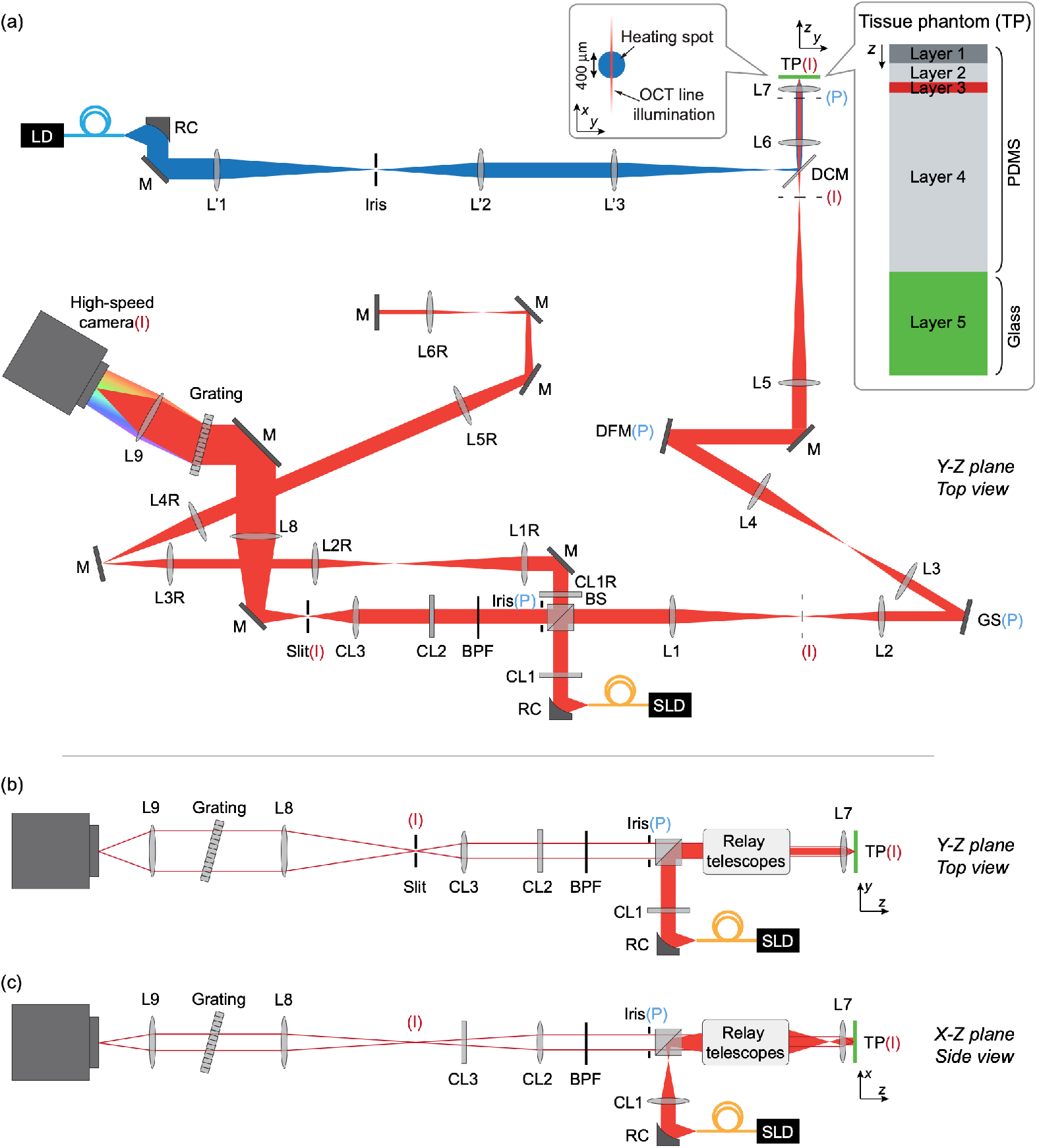
(a) Top-view of the optical setup including the line-field spectral-domain OCT (red path) and the laser heating (blue path). SLD: superluminescent diode, RC: reflective collimator, CL achromatic cylindrical lens doublet, L: achromatic lens doublet, BS: beam splitter (70T:30R), M: mirror, GS: galvo-scanning mirror, DFM: deformable mirror, DCM: (long-pass) dichroic mirror, TP: tissue phantom, LD: laser diode, BPF: band pass filter, P: pupil plane, I: image plane. The inset shows the overlap between the round heating beam spot and the OCT line illumination. The focal lengths of the lenses are: CL1 = CL1R = CL3 = 75 mm, L1 = L1R = L8 = 250 mm, L2 = L2R = 150 mm, L3 = L3R = 125 mm, L4 = L4R = CL2 = L2’ = 200 mm, L5 = L5R = 350 mm, L6 = L6R = L9 = 100 mm, L7 = 10 mm, L1’ = L3’ = 300 mm. The insets show the overlap between the heating beam and the OCT illumination field, and the tissue phantom assembly. (b) Simplified top view of the optical setup showing the illumination beam (solid beam) and two imaging rays in the detection path (red line). (c) Corresponding side view illustrating, with B, the anamorphic configuration based on cylindrical lenses (CL1–3).

The detection path, shown in two views in Fig. 1b,c follows an anamorphic imaging configuration by using two cylindrical lenses to allow independent control of magnification in the *x* and *y* dimensions, and thus independent control of the spatial and spectral resolutions. The OCT spectrometer is constructed using a 600 line/mm grating (Wasatch Photonics) and a high-speed camera (Phantom v641, 10-μm pixel size) for a resolution of 512 × 768 pixels (spatial × spectral). B-scans, with a static illumination field, are recorded for 500 ms with a frame rate of 10 kHz and 98-µs exposure times for a full imaging field of view of 2.3 mm laterally and 1.0 mm in depth. Flat glass windows are also added to the reference arm (not shown in schematic) for chromatic dispersion compensation between the reference and sample arms.

For laser heating, a blue diode beam (450-nm wavelength, 1-ms duration, 260-mW peak power) is coupled into a 200-µm multimode fiber. The fiber output is imaged onto the absorbing layer with a 400-µm diameter spot of uniform intensity. An iris is introduced at an intermediate image plane to sharpen the beam image edge and adjust the spot size. The beam is coupled into the OCT path using a long pass dichroic mirror and manually stirred so that the heating beam diameter overlaps with the line OCT illumination. The pulse duration and the delay for the OCT camera acquisition are controlled by a pulse/delay generator.

### 2.2. Tissue phantom fabrication and materials properties

A five-layer tissue phantom, emulating a simplified retina, was fabricated by spin coating several layers of poly(dimethysiloxane) (PDMS) to different thicknesses on top of a glass substrate. PDMS is a common tissue simulant material and has been used for retina phantom assemblies [23]. PDMS layers were fabricated using two-part, 10:1 mixing ratio, Sylgard 184 kits from Dow Corning, on top of a 1-inch borosilicate glass substrate (n°2 coverslip, ChemGlass) (Layer 5). The spin-coating recipes are given in Table 1. Layers were assembled in reverse layer label order, Layer 1, the top surface, being the last layer coated (see Fig. 1a inset). After spin-coating, samples were degassed in a desiccator for 30 minutes. All layers were then cured for 48 hours at room temperature before the next coating step. Curing at room temperature was chosen so not to degrade the dye, which visibly deteriorated at temperature above 40°C. Since the elastic modulus is dependent on the curing temperature [24], the same curing temperature was applied to all layers to simplify the mechanical model. To make the top, scattering PDMS layer (Layer 1), titanium particles (2.5wt%, 21-nm nominal diameter, Sigma Aldrich) were added to the base PDMS, previously diluted in hexane (3:1 weight ratio 3:1) to reduce the viscosity and hence facilitate the nano-powder mixing. The hexane was left to fully evaporate during stirring and regular sonication of the mixture for 24 hours prior to mixing with the curing agent and spin-coating. Layer 3, which aims to simulate the thin scattering and absorbing RPE layer, was fabricated using a PDMS/dye/titanium oxide mixture. First a saturated solution of hexane with Oil Red O dye, strong absorber at 450 nm, was prepared. The solution was then mixed with the PDMS base (1:1 weight ratio) to obtain a low viscosity mixture. Lastly, the titanium oxide nano-powder was added (2.5 wt% with respect to PDMS base) to the mixture, which was left under stirring and regular sonication for 24 hours. The mixture was then spin-coated at high speed.

**Table 1.**
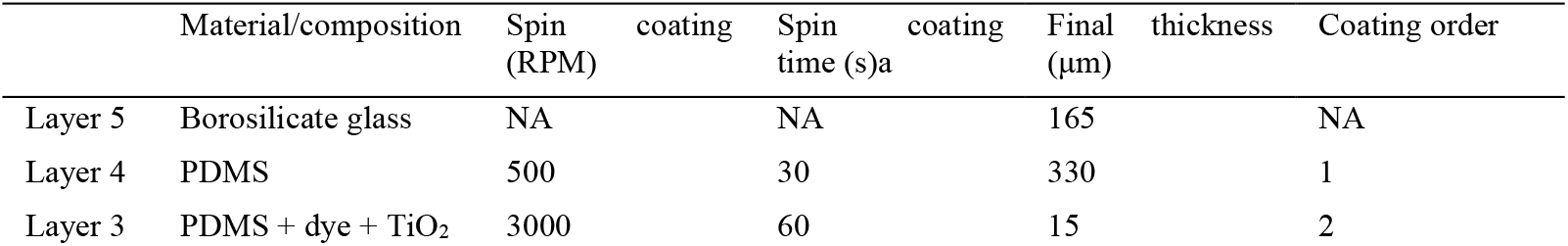
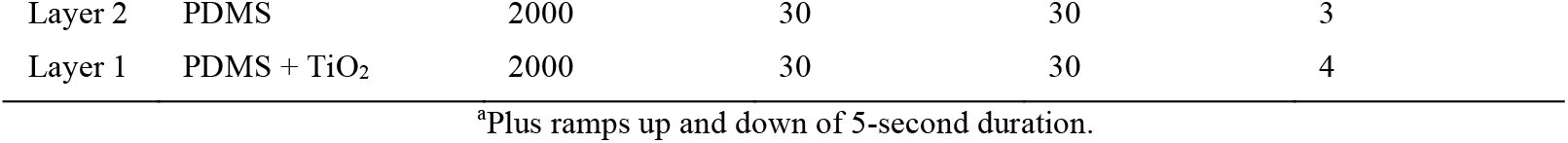
Tissue phantom layers and spin-coating recipes.

To measure the absorption coefficient of the dye-doped PDMS layer, a similar mixture but without the titanium particles (to avoid scattering), was prepared and spin-coated onto a glass substrate following the same recipe as for Layer 3. Light absorption in this layer, measured using the same 450-nm diode laser as in the heating experiments, was 8%. To calculate the absorption coefficient *μ*_*a*_, the thickness of that layer was measured by a profilometer. For simplicity the mechanical and thermal properties of all PDMS layers were considered identical. All material properties used in the modeling and references are listed in Table 2.

**Table 2.** Non-fitted material parameters used in model. *ρ:* density, *E:* Young’s modulus, *α*_*TE*_: coefficient of thermal expansion, *ν:* Poisson’s ratio, *κ*: coefficient of thermal conductivity, *c*_*p*_: specific heat capacity, *n*_*RT*_: refractive index at room temperature, *α*_*TE*_: thermo-optic coefficient, *μ*_*a*_: coefficient of optical absorption.

| | $\rho$ (kg.m<sup>-3</sup>) | $E$ (Pa) | $\alpha_{TE}$ (K<sup>-1</sup>)* | $\nu$ | $\kappa$ (W.m<sup>-1</sup>.K<sup>-1</sup>)* | $c_p$ (J.kg<sup>-1</sup>.K<sup>-1</sup>) | $n_{RT}$ | $\alpha_{TO}$ (K<sup>-1</sup>) | $\mu_a$ (mm<sup>-1</sup>)* (Layer 3 only) |
| --- | --- | --- | --- | --- | --- | --- | --- | --- | --- |
| PDMS | 970<sup>a</sup> | 1.3×10<sup>6</sup><br>[24] | 310×10<sup>-6</sup><br>[25] | 0.49<br>[25] | 0.17<sup>a</sup> | 1600<br>[26] | 1.41<br>[27] | -4.5×10<sup>-4</sup><br>[27] | 19.9 ± 0.8 |
| Borosilicate glass | 2230<sup>b</sup> | 63×10<sup>9</sup><sup>b</sup> | 3.3×10<sup>-6</sup><sup>b</sup> | 0.20<sup>b</sup> | 1.2<sup>b</sup> | 754<sup>c</sup> | NA | NA | NA |
| Air | 1.204<sup>c</sup> | NA | NA | NA | 25.87×10<sup>-3</sup><sup>c</sup> | 1006<sup>c</sup> | NA | NA | NA |

<sup>a</sup>Product datasheet (www.dow.com). <sup>b</sup>Product datasheet (www.schott.com). <sup>c</sup>www.engineeringtoolbox.com. Data taken at 23°C. References are indicated next to values.

### 2.3. OCT signal processing

OCT image reconstruction included the standard steps of background subtraction (of the sample and reference arms), wavenumber resampling, numerical dispersion compensation, and time referencing to cancel the local arbitrary phase [22]. In polar coordinates (*r,z*), the spatially static B-scan OCT signal can be described as a complex number:

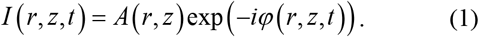

The phase difference Δ*φ* between two pixel coordinates (*r,z*)_1_ and (*r,z*)_2_ as a function of time is calculated by taking the argument of the product of the complex and complex conjugate numbers of the OCT signals at these locations:

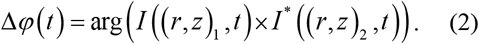

The OCT intensity image of the top 100 μm of the phantom is shown in Fig. 2a. The signal-to-noise ratio (SNR), which is directly related to phase sensitivity [28], is about 20-30 dB for the scattering layers of the tissue phantom. Because the top surface is reflective, while the other layers are scattering, it shows the highest SNR (maximum value ∼30 dB) and therefore has the lowest phase noise. We measured a minimum phase noise of 60 mrad at the top surface (*z* = 0) and at the center of the illumination beam (*r* = 0), where the OCT illumination intensity is the highest (center of the gaussian beam profile). The change in phase, Δ*φ*, can then be converted to change in single-pass optical path length, ΔOPL, by:

**Fig. 2.**
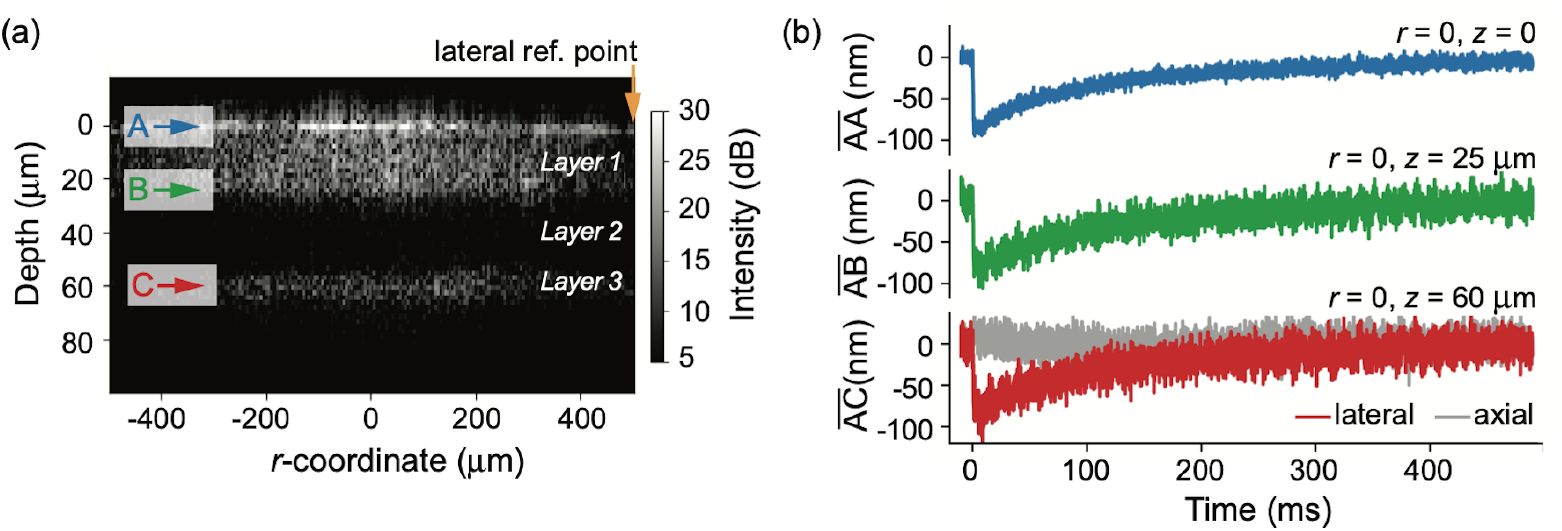
(a) B-scan OCT image showing the sample’s top layers. (b) Experimental changes in optical path lengths 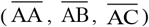 at the center of the heating beam (*r* = 0) for the three planes A, B, and C with corresponding depths *z* = 0, *z* = 25 μm, *z* = 60 μm. The planes are indicated by colored arrows in (a). The ΔOPLs were measured following heating by a single laser pulse at *t* = 0. The phase reference point (*r*_*ref*_ = 505 μm, *z* = 0) is indicated by a yellow arrow in (a). The low SNR signal that would be obtained for 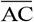 using an axial reference is shown in grey in C.

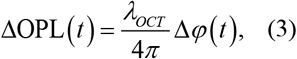

where *λ*_*OCT*_ is the central wavelength of the OCT illumination, yielding an OPL sensitivity of 4 nm at (*r* = 0, *z* = 0).

The *z*-coordinate of the top surface of the tissue phantom as a function of radius was found by tracking the peak OCT intensity averaged over 100 B-scan acquisitions, thus defining the plane A coordinates. Plane B and C were defined by translating plane A coordinates by 25 and 60 μm in depth, respectively.

### 2.4. Governing equations of the axisymmetric thermo-mechanical problem

The governing partial differential equations of equilibrium for an elastic medium in cylindrical coordinates can be expressed as [29]:

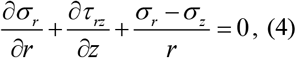

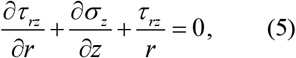

where *σ* _*r*_, *σ* _*z*_, are the normal stress components in the *r* and *z* directions and *τ* _*rz*_ is the shear stress in the *r-z* plane. The constitutive equations for an isotropic thermo-elastic medium are:

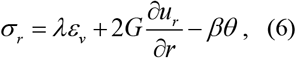

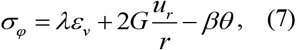

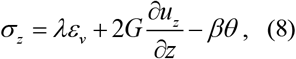

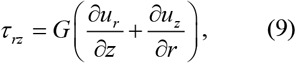

where *G* = *E* (2 (1−*ν*)) is the shear modulus with *E* the Young’s modulus and *ν* the Poisson’s ratio; *λ* = 2*Gν*/(1− 2*ν*) is the Lamé’s first parameter, *ε*_*v*_ = ∂*u*_*r*_/∂*r* + *u*_*r*_*/r* + ∂*u*_*z*_ /∂*z* is the volumetric strain. The term *βθ* represents the thermal stress in which *β* = 2*Gα*_*TE*_ (1+*ν*) (1− 2*ν*) is the thermo-mechanical coupling parameter, and *θ* is the temperature rise. The heat flux follows the Fourier’s heat conduction law:

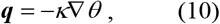

where ***q*** is the heat flux vector ***q*** = [*q*_*r*_, *q*_*z*_]^*T*^, *κ* is the coefficient of thermal conductivity and ∇ is the gradient operator. The heat flow in the *z* direction integrated over the time *t* is then:

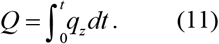

The heat diffusion equation can then be expressed as:

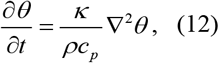

where *ρ* is the material density and *c*_*p*_ is the material specific heat capacity.

The solution to these equations, considering the initial and boundary conditions is described in the Supplemental Document. The external heat source flux *q*_*e*_ in the absorbing layer (layer 3) was modeled using the Beer-Lambert law, where the heat flux is:

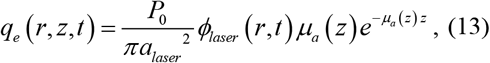

where *P*_*0*_, the incident laser power on layer 3, *a*_*laser*_ is the radius of the top-hat intensity profile, *ϕ*_*laser*_ is the spatio-temporal profile of the laser beam described below and *μ*_*a*_ is the optical absorption coefficient. *P*_*0*_ was calculated by taking into account the transmission through layer 1, which is scattering. The transmission of layer 1 was measured with the same diode laser used for heating and a power meter by comparing the transmitted power through an assembly comprising layers 1 to 5 (the tissue model) and a second assembly comprising layers 2 to 5 only to isolate the contribution from layer 1. Transmission of layer 1 is 50.0±0.5%.

The beam spatio-temporal profile was modeled as a square pulse in time and a square top-hat profile in space convolved with a Gaussian kernel to simulate blur and defocus [30]:

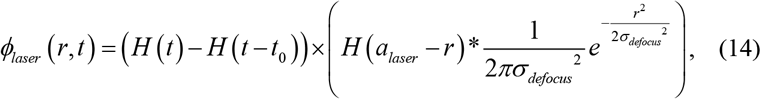

where *H* is the Heaviside function, *t*_*0*_ is the pulse duration (*t*_*0*_ = 1 ms), and *σ* _*defocus*_ is the Gaussian kernel width. For simplicity, we converted the volumetric heat source to a surface heat source of the following form:

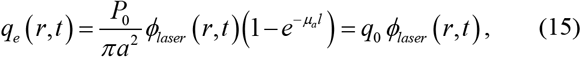

where 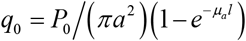 is the deposited heat flux amplitude and *l* is the thickness of layer 3. Given the diffusivity of PDMS, the volumetric heat source and the surface heat source become equivalent after the characteristic time of diffusion over the thickness of layer 3. This characteristic time is about 2 ms so fitting of the ΔOPL data after 2 ms using a surface heat source is justified and material parameters can be accurately fitted with this simplification. It is important to keep in mind that the calculated temperature at the absorbing layer differs whether a volumetric or surface heat source is used. Modeling a volumetric heat source is possible using the stiffness matrix method [31] but at the expense of longer computation time, which is acceptable after material properties are fitted. Using a volumetric or a surface heat source does not, however, affect temperature precision, which depends on the uncertainty of the fitted values. Finally, the external heat flow can be obtained by integrating the heat flux over time from 0 to *t*:

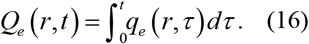

### 2.4. Arrhenius integral for determination of the therapeutic window

Thermal damage in retinal cells during transient hyperthermia have been approximated following the Arrhenius model, which assumes a first-order kinetics in denaturation of the critical molecular component, the ‘weak link’. Its concentration *D(t)* decreases at the rate [8]:

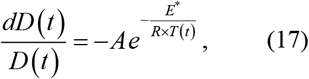

where *A* is the rate constant, *E\** is the activation energy, *R* is the gas constant (8.314 J.K^-1^.mol^- 1^) and *T* is the temperature as a function of time. The total decrease of *D* relative to its initial value *D*_*0*_ over the course of hyperthermia with duration *t*_*h*_ can be expressed by the Arrhenius integral Ω :

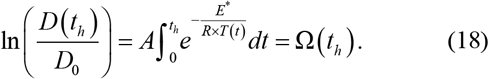

Values for *A* (1.6×10^55^ s^-1^) and *E\** (340 kJ/mol) have been calibrated and characterized in previous studies so that Ω = 1 corresponds to the RPE damage threshold [8]. No cellular response was observed for Ω < 0.1 thus the non-damaging therapeutic window was defined as 0.1 < Ω < 1 [7]. The Arrhenius integral can be directly calculated using the temperature course at any chosen point in the tissue.

## 3. Results

### 3.1. High-speed imaging of laser heating

The phase difference can be measured between any arbitrary pair of points. However, because we are interested in measuring the absolute vertical displacements of the layers, it is prudent to use a reference point whose phase remains largely unchanged following laser heating. Here, we have chosen a point at the surface of the sample (plane A, *z* = 0) far enough (*r*_*ref*_ = 505 μm) laterally to be least affected by the thermal expansion while maintaining a high SNR (Fig. 2a). Selection of such a reference point is possible since, with the line-field OCT, we have simultaneous access to the phase at all radial positions in a common-path configuration (in the sample arm only) [32]. In contrast, point-scan OCT only allows common-path phase referencing along the z axis (axial referencing) at a single radial position.

The complex values of the OCT signal at the three planes were averaged over 3 pixels in depth around the plane *z*-coordinates (± 1 pixel) before computing the phase values *φ* (*r, z*_*A*_, *t*), *φ* (*r, z*_*B*_, *t*), *φ* (*r, z*_*C*_, *t*). The phase reference *φ* (*r*_*ref*_, *z*_*A*_, *t*) was obtained by averaging the complex values of the OCT signal at the plane A *z*-coordinate and a radius offset of *r*_*ref*_ = 505 µm (± 1 pixel in *r* and *z*). The phase at *r*_*ref*_= −505 µm was also calculated to retrieve and cancel the surface tilt angle in the *r*-*z* plane.

The optical path difference was tracked as a function of time for the three planes of interest: the free surface (plane A, *z*_*A*_ = 0), an intermediate plane in layer 1 (plane B, *z*_*B*_ = 25 μm), and at the absorbing layer (plane C, *z*_*C*_ = 60 μm). For those planes, the ΔOPLs can be expressed as:

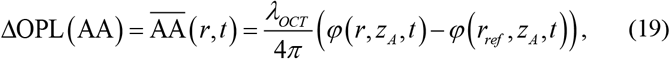

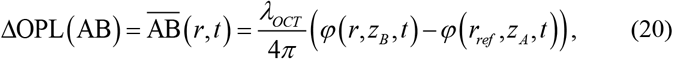

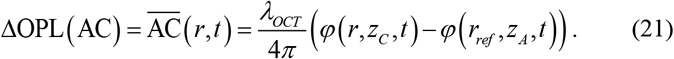

Figure 2b shows the 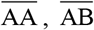, and 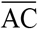 at the center of the heating beam (*r* = 0). Assuming for simplicity of interpretation that the reference phase is unchanged, 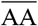 is directly opposite to the top surface displacement. Upon laser heating, the temperature of layer 3 increases and heat diffuses to adjacent layers. Heating leads to thermal expansion, so the layers deform and become thicker. It causes the top layer to move up (toward negative *z*) and 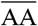 to decrease by the vertical displacement of the plane A (*u*_*z*_ (*r, z*_*A*_ = 0)). Meanwhile, as the temperature increases by *θ*, the index of refraction of PDMS decreases by Δ*n*, with Δ*n* = *α*_*TO*_*θ*, where *α*_*TO*_, the thermo-optic coefficient of PDMS, is negative. Therefore, as plane B moves up, the optical path 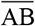 decreases by the additive effects of a shorter physical distance and a lower index of refraction. Since plane A also moves up and air is replaced by the polymer over the displacement thickness, the optical path 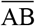 decrease is slightly reduced by this effect. Same goes for 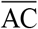. Because the thermal expansion (*α*_*TE*_) and thermo-optic (*α*_*TO*_) coefficients are intimately related, their combined effects resulted in 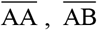, and 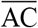 being of similar amplitude (Fig. 2b). The simplified expressions of 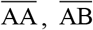, and 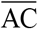 are:

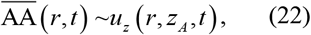

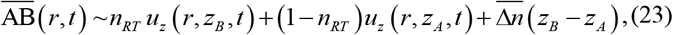

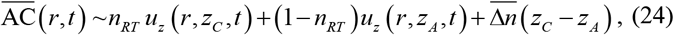

where *u*_*z*_ is normal displacement components in the *z* direction, *n*_*RT*_ is the index of refraction of PDMS at room temperature, and 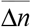 is the average change in refractive index induced by heating between the planes of interest. The exact expression and the derivation of these ΔOPLs can be found in the Supplemental Document (equations S2 and S7). For comparison, the change in optical path length that would be measured between the planes A and C using axial referencing, i.e. the phase difference between the planes at the same radial coordinate, would be:

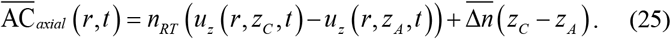

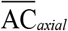 is shown along with 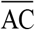 in Fig. 2b. It is worth noting that the SNR of this signal is much lower (close to zero) than the SNR of 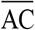 as the two terms in equation 10 compensate each other, making this signal hard to use.

### 3.2. Thermo-elastic modeling of transient heating

To model the ΔOPLs, one needs to calculate the vertical displacements *u*_*z*_ and distribution of the temperature change *θ* between the planes to obtain the change in refractive index Δ*n*. The displacements and temperature fields can be calculated by solving the thermo-elasticity problem. We followed the state space approach applied to layered media, introduced by Bufler [33] and Bahar [34]. This approach consists in reducing the partial differential equations (PDE) of coupled thermo-elasticity using integral transforms to a state space equation, taking advantage of the planar layered structure. The relationship between states vectors is then described by a transfer matrix [34,35] or a stiffness matrix [36] obtained by solving the state space equation. This method has been extensively used in macro-scale multi-layer problems in geophysics [37,38], with applications to seismology and nuclear waste management, but never at the microscale for biomaterials. The thermo-elastic problem can be analytically solved, in the transformed domain, for planar layers in a semi-infinite half space, which makes it particularly appropriate for the retina, as long as we assume a heat source radius being relatively small compared to the retina radius of curvature. In comparison, finite element methods (FEM) treat semi-infinite space by significantly extending the computational domain, therefore increasing the computation time. Moreover, for an axisymmetric configuration, inversing the solution from the transformed domain requires integration over two dimensions (the radial and temporal dimensions), whereas FEM perform integrations over three dimensions (the radial, axial, and temporal dimensions), which increases the computation time. The increased computation efficiency of the state space approach has been discussed in Refs [39,40] suggesting a >10× acceleration. Validation of the method and efficiency considerations can be found in the Supplemental Document (Stiffness matrix method (SMM) validation section). Rapid computing of the solution is critical for fitting convergence, fitting quality evaluation via bootstrapping, and precision assessment via Monte-Carlo simulations, all of which involve calculations of many solutions.

We followed the formulation by Ai et al. [31] to derive the state vector 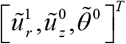 from the elastic equations, the generalized thermo-elastic Hooke Law, and the semi-coupled heat diffusion equation, where 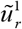 and 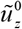 are the 1^st^ and 0^th^-order Hankel-Laplace (HL) transforms of the normal displacement components in the *r* and *z* directions, and 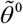 is the 0^th^-order HL transform of the temperature change. Numerical integration was then performed to find *u*_*r*_, *u*_*z*_, and *θ*, and the ΔOPLs were calculated using equations S2 and S7. The derivation of the state vector and the inverse numerical transform procedures are described in the Supplemental Document (Transfer and stiffness matrix derivation section).

### 3.3. Retrieval of the material properties

We assumed that the properties of the glass substrate are well known and, given the separation between the top of the assembly and the glass, that the contribution from the glass to the response was very small. We considered for simplicity that all PDMS layers have the same isotropic thermal and mechanical properties, including the layers containing titanium oxide particles, given their low weight fraction (2.5 wt%). The density *ρ*, the Poisson’s ratio *ν*, and the specific heat capacity *c*_*p*_ are all well characterized for PDMS [25,26] and are considered fixed parameters. Similarly, we assumed the index of refraction *n* at room temperature (23°C) and the thermo-optic coefficient *α*_*TO*_ [27] to be known. On the other hand, the coefficient of thermal expansion *α*_*TE*_ strongly depends on the curing temperature, which leads to uncertainty in its value [25], and a range of coefficient of thermal conductivity *κ* can be found in the literature. We therefore considered *α*_*TE*_ and *κ* as fitting parameters. Although the Young’s modulus *E* depends on the curing temperature [24], it has little effect on the state vector (see Supplemental Document, Elastic modulus effects section). Lastly, the absorption coefficient *μ*_*a*_ was fitted through the deposited heat magnitude *q*_0_ with 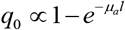, where *l* is the absorbing layer thickness. We further assumed that these properties are temperature independent, given the small temperature range of the study (<10°C). We accounted for the potential defocus of the heating laser beam by adding a Gaussian blur to the ideal disk (top-hat) intensity profile (equation 14) [30], which added two geometrical fitting parameters to the model (width of the blur *σ*_*defocus*_ and top-hat profile radius *a*_*laser*_). This brought the total number of fitting parameters to five: *α*_*TE*_, *q*_*0*_, *κ, σ*_*defocus*_, *and a*_*laser*_.

Fitting of the parameters was performed in two steps. First, these properties were fitted by minimizing the sum of the squares of the residuals between the model and the experimental data 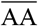, which showed the highest SNR, using the exact ΔOPL expressions. The laser geometrical parameters, *σ*_*defocus*_ *and a*_*laser*_, were mainly fitted through the spatial shape of the response, and the coefficient of thermal conductivity was mainly fitted through the temporal decay of the response. Importantly, however, 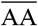 scales linearly with both *α*_*TE*_ and *q*_*0*_ so fitting of 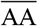 did not allow decorrelation between *α*_*TE*_ and *q*_*0*_, therefore a product *α*_*TE*_ × *q*_0_ was fitted through the magnitude of the response. Second, to find *α*_*TE*_ and *q*_*0*_, we used the experimental data 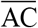, which is only proportional to *q*_*0*_. The fitted parameters found in the first step were used to solve for 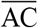, and then amplitude of 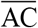 was fitted to determine the absolute values of *α*_*TE*_ and *q*_*0*_, as described in the Supplemental Document (Material property fitting procedure section). The comparison between the fitted model and the data is shown in Fig. 3 for 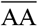 and 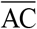. The data are well reproduced in both the spatial (radial) and temporal dimensions, thus suggesting high fidelity of the model. The uniformity of the residuals (see Supplemental Document) over the entire data set also attests for the accuracy of the model and the absence of systematic errors. The resulting fitted values are listed in Table 3. We note a good agreement between the fitted and initial values for *α*_*TE*_ (286 vs 310 K^-1^), *κ* (0.175 vs 0.17 W.m^-1^.K^-1^), and *μ*_*a*_ (20.3 mm^-1^ vs 19.9 mm^-1^). The uncertainties in parameter determination were obtained by fitting bootstrapped data, resampled over a total of 200 iterations, and are listed in Table 3. The obtained uncertainties are an order of magnitude lower than for the starting values of *α*_*TE*_ and *κ*, hence demonstrating the high precision of the parameter determination using this method. We also note that the obtained precision on *μ*_*a*_ (2.5%) is much higher than the required accuracy for therapeutic titration (20%). We finally verified the low correlation between the parameters (see Parameter correlations section in Supplemental Document) obtained by repeated fitting of the bootstrapped data.

**Table 3.** Non-fitted and fitted PMDS material parameters. *α*_*TE*_: coefficient of thermal expansion, *κ*: coefficient of thermal conductivity, *μ*_*a*_: coefficient of optical absorption.

| | $\alpha_{TE}$ ($K^{-1}$) | $\kappa$ ($W \cdot m^{-1} \cdot K^{-1}$) | $\mu_a$ ($mm^{-1}$) (Layer 3 only) |
| --- | --- | --- | --- |
| PDMS (not fitted) | $310 \times 10^{-6}$ [25] | 0.17<sup>a</sup> | $19.9 \pm 0.8$ |
| PDMS (fitted) | $286 \times 10^{-6} \pm 3 \times 10^{-6}$ | $0.175 \pm 0.03$ | $20.3 \pm 0.5$ |

<sup>a</sup>Product datasheet (www.dow.com). References are indicated next to values.

**Fig. 3.**
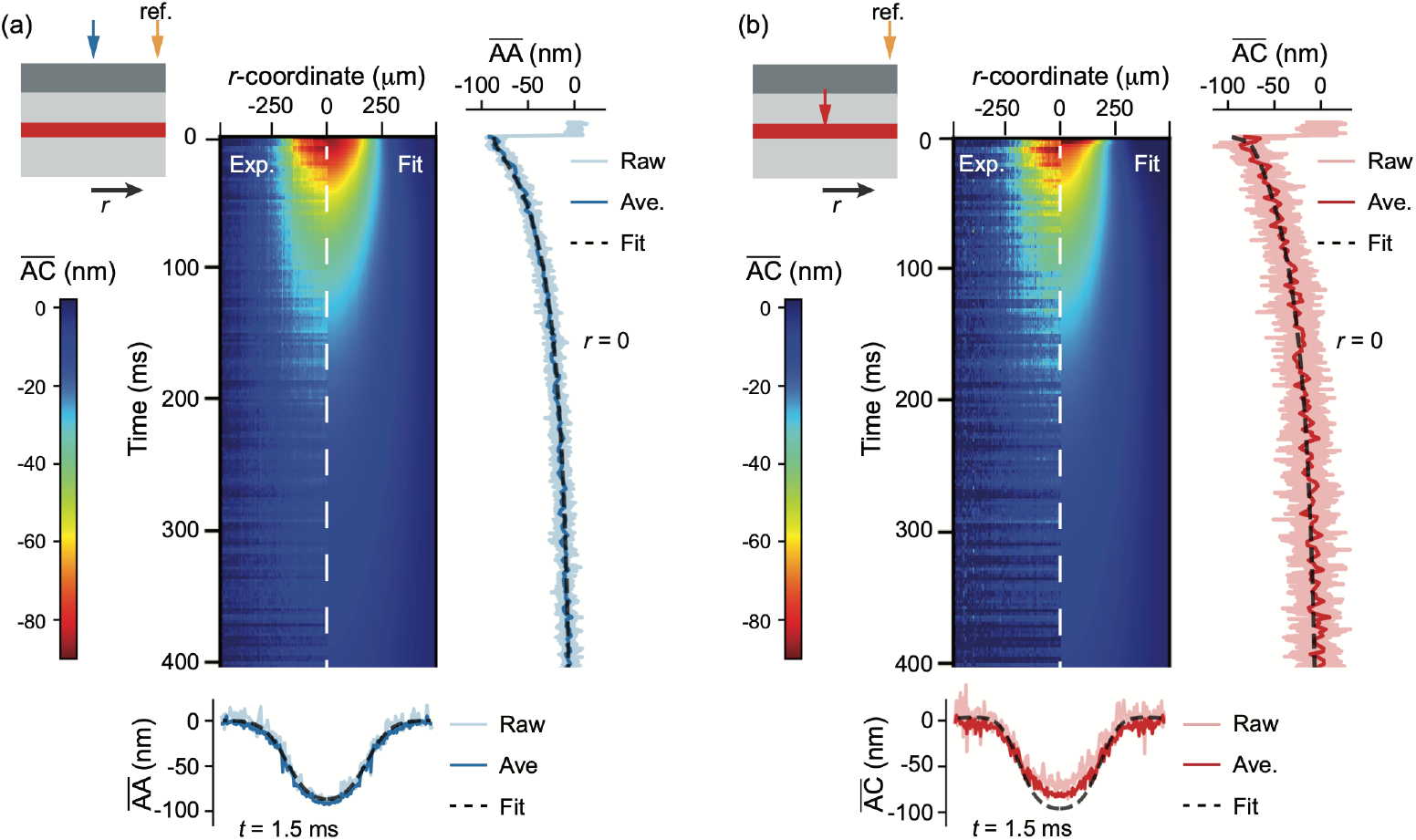
(a) 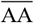 and (b) 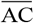 experimental data (left halves) compared to the fitted model (right halves) in space (radial) and time. The temporal profile of the response at the center of the heating beam (*r* = 0) and its radial profile shortly after the heating pulse was turned off (*t* = 1.5 ms) are shown on the right and below the maps. The model was fitted on the rolling average of the raw experimental data.

### 3.4. Temperature estimation and precision

Since we are ultimately interested in the temperature, the best-fit parameters can be used to compute the temperature increase *θ* at any point in space and time. For non-damaging retinal therapy, the temperature rise at the RPE, represented here by the absorbing PDMS layer, is of utmost importance. Figure 4a shows the temporal course of the temperature at that layer. Temperature increased during the 1 ms laser pulse, and then decreased slowly due to heat diffusion. Figure 4b shows two snapshots of the temperature distribution at the end of the heating pulse (*t* = 1 ms), when heat is relatively confined, and after 4 ms, when heat has diffused radially and axially. To estimate the temperature precision of the calculations, we ran Monte-Carlo simulations using the fitted parameters with their uncertainty over 1000 iterations to compute the peak temperature (*t* = 1 ms) at the center of the beam at the absorbing layer (*r* = 0, *z*_*C*_ = 60 μm), and found a temperature uncertainty of ±0.25°C. These simulations at a single point in time and space took about 20 seconds. For comparison, running the same number of iterations using the COMSOL model would take approximately 6 hours.

**Fig. 4.**
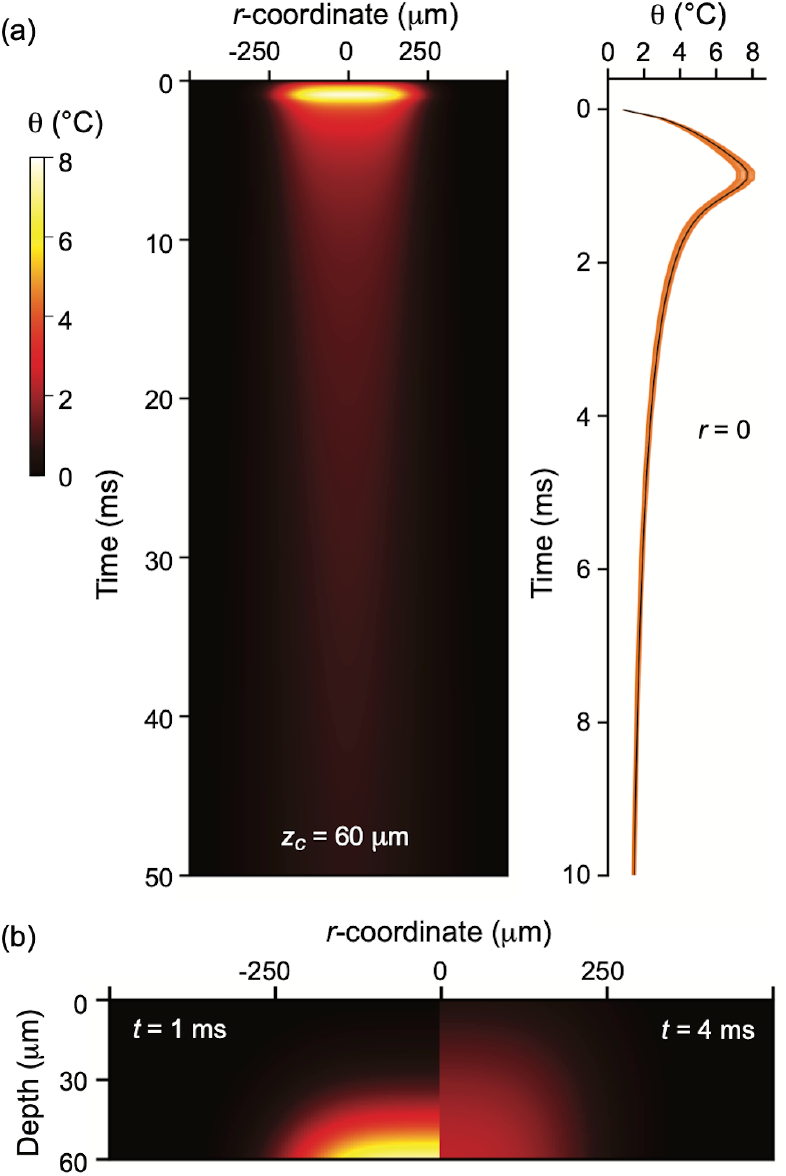
(a) Radial and temporal distribution of temperature at the absorbing layer (*z*_*C*_ = 60 μm). The temporal course at the center of the heating beam (*r* = 0) calculated with the best-fit parameters (black line) is shown on the right. Monte-Carlo simulations (*n* = 50) considering parameter uncertainties are shown in orange to illustrate the uncertainty in the temperature estimation. (b) Spatial distributions of temperature at 1 and 4 ms, showing heat diffusion into the sample.

## 4. Discussion

Determination of the temporal and spatial distribution of the temperature rise in tissue is critical for evaluating the safe therapeutic window and the extent of potential thermal damage. Both are typically assessed using the Arrhenius equation, which describes the first order kinetics of the thermal denaturation. The therapeutic window for non-damaging hyperthermia corresponds to the Arrhenius integral value between 0.1 and 1, where values above 1 correspond to permanent damage. With the 2.5% precision of the absorption coefficient determination in our approach, the uncertainty in Ω is about 0.17, which is well within the therapeutic window. Uncertainty of 0.50 in Arrhenius integral, approaching the boundary of the therapeutic window corresponds to 2.5 times lower precision in determination of the absorption coefficient (i.e., 6.25%). At the current laser settings, this would translate to a temperature rise uncertainty of about 0.65°C. Precision of this method exceeds the other methodologies for temperature-controlled retinal treatment and is more than sufficient for titration the non-damaging thermal therapy. In the current example, the absorption coefficient determination and the temperature calculations are performed way below the damage threshold (Ω < 10^−3^), which enables repeated measurements (multiple heating pulses) for further data averaging.

We are working on adapting this approach to *in-vivo* measurements and hope to achieve a similar level of the temperature precision despite the potential difficulties with tissue movements. Here, with the tissue phantom, we only considered the phase information from two planes (A and C), as they enable sufficient precision in determination of the material parameters. However, more data points from additional planes (e.g., plane B) are available for fitting, which would increase the precision and allow additional fitting parameters. On one hand, retina offers many scattering planes from which phase can be extracted, while on the other hand, we expect a larger number of variables for the retina. It remains to be seen whether the increased number of experimental data points provided by the additional layers will compensate for the increased number of unknown parameters.

Retinal thermal expansion coefficient is often assumed to be equal to that of water [17], but a recent study suggested otherwise and estimated it to be four times larger [16]. We can also expect this coefficient to be anisotropic (albeit transverse isotropic) for the retina, given its anisotropic cellular structure. The same might apply to the coefficient of thermal conductivity and, for the same reason, layers could exhibit different Poisson’s ratios. Conversely, the index of refraction of the retina is relatively uniform within 1% across the layers and has been reasonably determined [41]. Questions remain, however, regarding the thermo-optic coefficient (*α*_*TO*_), which might more strongly depend on the solid content [42,43]. It also has been shown that the elastic modulus varies throughout the retina [19], and is two-to-three orders of magnitude lower than for PDMS (kPa instead of MPa). We therefore expect it to play a more significant role than in the tissue phantom. In addition, the present model assumes a uniform laser irradiance and a uniform absorption coefficient within the laser spot. For *in-vivo* measurements, we can ignore potential cellular-scale variations (∼ 30 μm) in the absorption coefficient and laser-intensity variations (∼ 10 μm) in the heat source profile. since the variations would be homogenized within 1 ms of the laser pulse, as heat diffuses by about 30 μm over that duration.

## 5. Conclusion

We present a methodology for determination of the optical and thermal parameters in multi-layered materials, applicable to retinal photothermal therapies. We demonstrated that, by matching the optical path length changes, recorded with line-field phase-resolved optical coherence tomography, across the beam width and along the full course of heating and cooling, one can precisely identify the material properties. Based on model fittings, we estimated the temperature with a precision of about 0.2°C for a heating amplitude not exceeding 8°C during 1 ms—well within the safety range of the retina [5]. *In-situ* measurements of the retinal heating in every patient should allow precise titration of the energy deposition for photocoagulation and non-damaging laser therapy. We finally note that the present methodology is also generally applicable to any planar multi-layered assembly, as long as it provides an OCT signal with sufficiently high SNR. Beyond biomaterials and tissues, the method could be extended, for example, to coating and surface metrology [44] or artwork studies [45].

## Acknowledgements

We thank Vimal Pandiyan, Ramkumar Sabesan, Fabio Feroldi, and Austin Roorda for discussions regarding the optical design. DV thanks Ludwig Galambos for profilometry measurements, Zhi Yong Ai for insights on the transfer and stiffness matrix method, and Mohajeet Bhuckory for discussions on the translation to *in-vivo* measurements.

## Data availability

All study data are included in the article and/or in the Supplemental Document.

## Supplemental document

See Supplement 1 for supporting content.

