## Supplemental Document 1 for "Interferometric imaging of thermal expansion for temperature control in retinal laser therapy"

### 1. Optical path lengths and lateral versus axial phase reference

The phase and optical path changes following thermal perturbation using a lateral reference point can be derived using a virtual undisturbed plane (plane V, see Fig. S1). The single-pass ΔOPL $\overline{\mathrm{AA}}$ is therefore the difference in optical paths between the plane A at radius $r$ and the virtual plane (blue path in Fig. S1) and the plane A at radius $r_{ref}$ and the virtual plane (yellow path in Fig. S1):

$$\overline{\mathrm{AA}}(r,t)=\int_{z_V}^{z_A+u_z(r,z_A,t)} n_{air}(r,z)\,dz-\int_{z_V}^{z_A+u_z(r_{ref},z_A,t)} n_{air}(r_{ref},z)\,dz, \quad \text{(S1)}$$

where $z_A$ and $z_V$ are the $z$-coordinates of the planes A and V, $u_z$ is the vertical displacement (here of the plane A), $n$ is the index of refraction of the materials (here $n_{air}$) along the integration paths. Assuming negligible heating of the air above the tissue phantom, $n_{air}$ remains unchanged and equal to 1. $\overline{\mathrm{AA}}$ can be simplified as:

$$\overline{\mathrm{AA}}(r,t)=z_A+u_z(r,z_A,t)-z_V-z_A-u_z(r_{ref},z_A,t)+z_V=u_z(r,z_A,t)-u_z(r_{ref},z_A,t). \quad \text{(S2)}$$

Further assuming negligible surface displacement at radius $r_{ref}$ gives:

$$\overline{\mathrm{AA}}(r,t)\sim u_z(r,z_A,t). \quad \text{(S3)}$$

This additional simplification step is not taken during simulations of the optical paths.

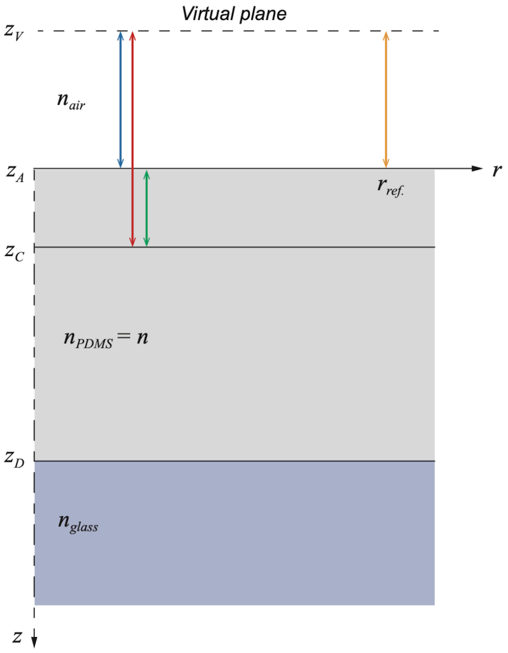


Fig. S1. OPL integration paths used to calculated $\overline{\mathrm{AA}}$ (blue and yellow paths), $\overline{\mathrm{AC}}$ (red and yellow paths), and $\overline{\mathrm{AC}}_{axial}$ (green path).

Similarly, the $\overline{\mathrm{AC}}$ can be derived by subtracting the optical paths between the planes C and V at radius *r* (red path in Fig. S1) and the planes A and V at radius $r_{ref}$ (yellow path in Fig. S1):

$$\overline{\mathrm{AC}}(r,t) = \int_{z_V}^{z_C + u_z(r,z_C,t)} n(r,z)\,dz - \int_{z_V}^{z_A + u_z(r_{ref},z_A,t)} n_{air}(r_{ref},z)\,dz\,. \quad \text{(S4)}$$

The red path is split into two paths comprising the air path and the PDMS path, giving:

$$\overline{\mathrm{AC}}(r,t) = \int_{z_V}^{z_A + u_z(r,z_A,t)} n_{air}(r,z)\,dz + \int_{z_A + u_z(r,z_A,t)}^{z_C + u_z(r,z_C,t)} n_{PDMS}(r,z,T(t))\,dz - \int_{z_V}^{z_A + u_z(r_{ref},z_A,t)} n_{air}(r_{ref},z)\,dz\,. \quad \text{(S5)}$$

Along the integration path in PDMS, the index of refraction of PDMS ( $n_{PDMS}$ simplified as $n$ ) changes, because of the temperature increase $\theta$, from $n(r,z,23°\mathrm{C}) = n_{RT}$ (room-temperature value) to $n_{RT} + \Delta n(r,z,\theta(t))$ where $\Delta n(r,z,\theta(t)) = \alpha_{TO}\theta$ and $\alpha_{TO}$ is the thermo-optic coefficient. Integrating the index of refraction is equivalent to averaging its value over the integration path, then equation S5 becomes:

$$\overline{\mathrm{AC}}(r,t) = u_z(r,z_A,t) - u_z(r_{ref},z_A,t) + \left(n_{RT} + \overline{\Delta n}\right)\left(z_C - z_A + u_z(r,z_C,t) - u_z(r,z_A,t)\right), \quad \text{(S6)}$$

where $\overline{\Delta n}$ is the average value of $\Delta n$ between the planes A and C. The terms in $\overline{\Delta n} \times u_z$ are negligible, yielding:

$$\overline{\mathrm{AC}}(r,t) = u_z(r,z_A,t) - u_z(r_{ref},z_A,t) + n_{RT}\left(u_z(r,z_C,t) - u_z(r,z_A,t)\right) + \overline{\Delta n}\left(z_C - z_A\right) + n_{RT}\left(z_C - z_A\right)\,. \quad \text{(S7)}$$

Again, if we consider small displacements at the radius $r_{ref}$, $\overline{\mathrm{AC}}$ can be approximated as:

$$\overline{\mathrm{AC}}(r,t) \sim n_{RT}\,u_z(r,z_C,t)+(1-n_{RT})u_z(r,z_A,t)+\overline{\Delta n}(z_C-z_A). \text{ (S8)}$$

The last term $n_{RT}(z_C-z_A)$ is a constant (initial ΔOPL) and can be dropped as we are only interested in OPL changes. For $\overline{\mathrm{AB}}$, C indices are replaced by B indices.

$$\overline{\mathrm{AB}}(r,t) \sim n_{RT}\,u_z(r,z_B,t)+(1-n_{RT})u_z(r,z_A,t)+\overline{\Delta n}(z_B-z_A). \text{ (S9)}$$

In contrast, the axial phase difference between the planes A and C (green path in Fig. S1), which is calculated at the same radius and does not consider a lateral reference point, is:

$$\overline{\mathrm{AC}}_{axial}(r,t)=\int_{z_A+u_z(r,z_A,t)}^{z_C+u_z(r,z_C,t)} n_{PDMS}(r,z)\,dz, \qquad \text{(S10)}$$

$$\overline{\mathrm{AC}}_{axial}(r,t)=n_{RT}\left(u_z(r,z_C,t)-u_z(r,z_A,t)\right)+\overline{\Delta n}(z_C-z_A)+n_{RT}(z_C-z_A) \qquad \text{(S11)}$$

$$\overline{\mathrm{AC}}_{axial}(r,t)=n_{RT}\left(u_z(r,z_C,t)-u_z(r,z_A,t)\right)+\overline{\Delta n}(z_C-z_A), \qquad \text{(S12)}$$

with the constant term dropped.

### 2. Transfer and stiffness matrix derivation

Here we reproduce the governing equations from the Materials and Methods section and further elaborate on the derivation to find the stiffness matrix and the state vector solutions. The governing partial differential equations (PDE) of equilibrium are [1]:

$$\frac{\partial\sigma_r}{\partial r}+\frac{\partial\tau_{rz}}{\partial z}+\frac{\sigma_r-\sigma_z}{r}=0, \text{(S13)}$$

$$\frac{\partial\tau_{rz}}{\partial r}+\frac{\partial\sigma_z}{\partial z}+\frac{\tau_{rz}}{r}=0, \qquad \text{(S14)}$$

where $\sigma_r$, $\sigma_z$, are the normal stress components in the *r* and *z* directions and $\tau_{rz}$ is the shear stress in the *r*-*z* plane. The constitutive equations for an isotropic thermo-elastic medium are:

$$\sigma_r=\lambda\varepsilon_v+2G\frac{\partial u_r}{\partial r}-\beta\theta, \quad \text{(S15)}$$

$$\sigma_\varphi=\lambda\varepsilon_v+2G\frac{u_r}{r}-\beta\theta, \quad \text{(S16)}$$

$$\sigma_z=\lambda\varepsilon_v+2G\frac{\partial u_z}{\partial z}-\beta\theta, \quad \text{(S17)}$$

$$\tau_{rz}=G\left(\frac{\partial u_r}{\partial z}+\frac{\partial u_z}{\partial r}\right), \qquad \text{(S18)}$$

where $G=E/\left(2(1-\nu)\right)$ is the shear modulus with $E$ the Young's modulus and $\nu$ the Poisson's ratio; $\lambda=2G\nu/(1-2\nu)$ is the Lamé's first parameter, $\varepsilon_v=\partial u_r/\partial r+u_r/r+\partial u_z/\partial z$ is the volumetric strain, $\beta\theta$ is the thermal stress in which $\beta=2G\alpha_{TE}(1+\nu)/(1-2\nu)$, and $\theta$

is the temperature rise. Substituting equation S15–18 into S13–14, we obtain the following PDE relating displacements and temperature:

$$\nabla^2 u_r + \frac{1}{1-2\nu}\frac{\partial \varepsilon_v}{\partial r} - \frac{1}{r^2}u_r - \frac{\beta}{G}\frac{\partial \theta}{\partial r} = 0, \qquad \text{(S19)}$$

$$\nabla^2 u_z + \frac{1}{1-2\nu}\frac{\partial \varepsilon_v}{\partial z} - \frac{\beta}{G}\frac{\partial \theta}{\partial z} = 0, \qquad \text{(S20)}$$

where $\nabla^2 = \partial^2/\partial r^2 + 1/r\,\partial/\partial r + \partial^2/\partial z^2$ is the Laplacian operator. We recall the heat flux expression, which follows the Fourier's heat conduction law:

$$\boldsymbol{q} = -\kappa \nabla \theta, \qquad \text{(S21)}$$

where $\boldsymbol{q} = [q_r, q_z]^T$ is the heat flux vector, $\kappa$ is the coefficient of thermal conductivity, and $\nabla$ is the gradient operator $\nabla = [\partial/\partial r, \partial/\partial z]^T$. The heat flow in the *z* direction integrated over the time *t* is:

$$Q = \int_0^t q_z dt. \qquad \text{(S22)}$$

The heat diffusion equation is:

$$\frac{\partial \theta}{\partial t} = \frac{\kappa}{\rho c_p}\nabla^2 \theta = a\nabla^2\theta, \qquad \text{(S23)}$$

where $\rho$ is the material density, $c_p$ is the material specific heat capacity, and $a = \kappa/(\rho c_p)$ is the coefficient of thermal diffusivity.

Equations S19–23 can be transformed into the Hankel-Laplace (HL) domain, assuming initially undisturbed media, which is required for Laplace transformation. The combined $m^{\text{th}}$-order Hankel and Laplace transforms of a function $f(r,z,t)$ is $\tilde{f}^m(\xi,z,s)$ given by [2]:

$$\tilde{f}^m(\xi,z,s) = \int_0^{+\infty}\int_0^{+\infty} f(r,z,t) J_m(\xi r) r e^{-st} dr dt, \qquad \text{(S24)}$$

where $\xi$ and $s$ are the Hankel and Laplace variables, respectively, and $J_m$ is the $m^{\text{th}}$-order Bessel function of the first kind. The inverse Hankel-Laplace transform is defined as:

$$f(r,z,t) = \frac{1}{2\pi i}\int_{c-i\infty}^{c+i\infty}\int_0^{+\infty} \tilde{f}^m(\xi,z,s) J_m(\xi r)\xi e^{st} d\xi ds. \qquad \text{(S25)}$$

Taking the $1^{\text{st}}$-order Hankel and Laplace transforms of equation S19 gives:

$$\frac{d^2\tilde{u}_r^1}{dz^2} - \frac{2(1-\nu)}{1-2\nu}\xi^2\tilde{u}_r^1 - \frac{1}{1-2\nu}\xi\frac{\partial \tilde{u}_z^0}{\partial z} + \frac{\beta}{G}\xi\tilde{\theta}^0 = 0, \qquad \text{(S26)}$$

and taking the $0^{\text{th}}$-order Hankel and Laplace transforms of equations S20 and S23 gives:

$$\frac{2(1-\nu)}{1-2\nu}\frac{d^2\tilde{u}_z^0}{dz^2} - \xi^2\tilde{u}_z^0 + \frac{1}{1-2\nu}\xi\frac{\partial \tilde{u}_r^1}{\partial z} - \frac{\beta}{G}\frac{\partial \tilde{\theta}^0}{\partial z} = 0, \qquad \text{(S27)}$$

$$s\tilde{\theta}^0 - a\frac{d^2\tilde{\theta}^0}{dz^2} + a\xi^2\tilde{\theta}^0 = 0. \qquad \text{(S28)}$$

Defining a vector $\mathbf{W}$ by:

$$\mathbf{W}=\left[\tilde{u}_r^1,\tilde{u}_z^0,\tilde{\theta}^0,\frac{d\tilde{u}_r^1}{dz},\frac{d\tilde{u}_z^0}{dz},\frac{d\tilde{\theta}^0}{dz}\right]^T=\left[\mathbf{\Lambda},\frac{d\mathbf{\Lambda}}{dz}\right]^T, \qquad \text{(S29)}$$

where $\mathbf{\Lambda}=\left[\tilde{u}_r^1,\tilde{u}_z^0,\tilde{\theta}^0\right]^T$, equations S26–28 can be combined to obtain:

$$\frac{d\mathbf{W}(\xi,z,s)}{dz}=\mathbf{\Phi W}(\xi,z,s), \qquad \text{(S30)}$$

with:

$$\mathbf{\Phi}=\begin{bmatrix} & 0_{3\times3} & & & \boldsymbol{I}_{3\times3} & \\ \frac{2(1-\nu)}{1-2\nu}\xi^2 & 0 & -\frac{\beta}{G}\xi & 0 & \frac{1}{1-2\nu}\xi & 0 \\ 0 & \frac{1-2\nu}{2(1-\nu)}\xi^2 & 0 & -\frac{1}{2(1-\nu)}\xi & 0 & \frac{1-2\nu}{2(1-\nu)}\frac{\beta}{G} \\ 0 & 0 & \frac{s}{a}+\xi^2 & 0 & 0 & 0 \end{bmatrix}. \qquad \text{(S31)}$$

$0_{3\times3}$ is a 3×3 zero matrix and $\boldsymbol{I}_{3\times3}$ is a 3×3 identity matrix. A general solution satisfying equation S30 can be written as:

$$\mathbf{W}(\xi,z,s)=\mathbf{X}(\xi,s)e^{-\lambda z}, \qquad \text{(S32)}$$

where $\mathbf{X}$ and $\lambda$ are to be determined, which involves solving the eigenvalue problem:

$$\lambda\mathbf{X}=\mathbf{\Phi X}. \qquad \text{(S33)}$$

The eigenvalues are $\lambda_1=\xi$, $\lambda_2=\xi$, $\lambda_3=-\xi$, $\lambda_4=-\xi$, $\lambda_5=\left(s/a+\xi^2\right)^{1/2}$, and $\lambda_6=-\left(s/a+\xi^2\right)^{1/2}$, and the corresponding solution can be written as:

$$\mathbf{W}(\xi,z,s)=(\boldsymbol{c}_1+\boldsymbol{c}_2z)e^{\xi z}+(\boldsymbol{c}_3+\boldsymbol{c}_4z)e^{-\xi z}+\boldsymbol{c}_5e^{\sqrt{\frac{s}{a}+\xi^2}z}+\boldsymbol{c}_6e^{-\sqrt{\frac{s}{a}+\xi^2}z}, \qquad \text{(S34)}$$

with:

$$\boldsymbol{c}_1=\boldsymbol{m}_1l_1+\boldsymbol{m}_2l_2, \qquad \text{(S35)}$$

$$\boldsymbol{c}_2=\boldsymbol{m}_1l_2, \qquad \text{(S36)}$$

$$\boldsymbol{c}_3=\boldsymbol{m}_3l_3+\boldsymbol{m}_4l_4, \qquad \text{(S37)}$$

$$\boldsymbol{c}_4=\boldsymbol{m}_3l_4, \qquad \text{(S38)}$$

$$\boldsymbol{c}_5=\boldsymbol{m}_5l_5, \qquad \text{(S39)}$$

$$\boldsymbol{c}_6=\boldsymbol{m}_6l_6, \qquad \text{(S40)}$$

where $\boldsymbol{m}_i$ can be derived from the eigenvectors of $\mathbf{\Phi}$ and $l_i$ are arbitrary constants. For concision the vectors are not reproduced here but they are trivially derived. Considering a single finite layer, the vectors $\mathbf{\Lambda}(\xi,z,s)$ at depth z and $\mathbf{\Lambda}(\xi,0,s)$ at $z = 0$ can be expressed in terms of $\mathbf{L}$ :

$$\begin{bmatrix}\mathbf{\Lambda}(\xi,0,s)\\ \mathbf{\Lambda}(\xi,z,s)\end{bmatrix}=\begin{bmatrix}\mathbf{M}(\xi,0,s)\\ \mathbf{M}(\xi,z,s)\end{bmatrix}\mathbf{L}, \qquad \text{(S41)}$$

with $\mathbf{L}=\begin{bmatrix}l_1 & l_2 & l_3 & l_4 & l_5 & l_6\end{bmatrix}^T$. $\mathbf{M}$ can be derived from S34–40. The stress vector $\mathbf{V}=\left[\tilde{\tau}^1_{rz},\tilde{\sigma}^0_z,\tilde{Q}^0\right]^T$ is related to $\mathbf{\Lambda}$ by the $\mathbf{S}$ matrix obtained by transforming equation S17, S18, S21, and S22 into the HL domain:

$$\tilde{\tau}^1_{rz}=G\left(\frac{\partial\tilde{u}^1_r}{\partial z}-\xi\tilde{u}^0_z\right), \qquad \text{(S42)}$$

$$\tilde{\sigma}^0_z=\frac{2G\nu}{1-2\nu}\xi\tilde{u}^1_r+2G\frac{1-\nu}{1-2\nu}\frac{\partial\tilde{u}^0_z}{\partial z}-\beta\tilde{\theta}^0, \qquad \text{(S43)}$$

$$\tilde{Q}^0=-\frac{\kappa}{s}\frac{\partial\tilde{\theta}^0}{\partial z}, \qquad \text{(S44)}$$

giving:

$$\mathbf{V}(\xi,z,s)=\mathbf{S}(\xi,s)\mathbf{W}(\xi,z,s), \qquad \text{(S45)}$$

$$\mathbf{S}(\xi,s)=\begin{bmatrix}0 & -G\xi & 0 & G & 0 & 0\\ \frac{2G\nu}{1-2\nu}\xi & 0 & -\beta & 0 & 2G\frac{1-\nu}{1-2\nu} & 0\\ 0 & 0 & 0 & 0 & 0 & -\frac{\kappa}{s}\end{bmatrix}. \qquad \text{(S46)}$$

The vectors $\mathbf{V}(\xi,z,s)$ and $\mathbf{V}(\xi,0,s)$ can also be expressed in terms of $\mathbf{L}$ :

$$\begin{bmatrix}-\mathbf{V}(\xi,0,s)\\ \mathbf{V}(\xi,z,s)\end{bmatrix}=\begin{bmatrix}\mathbf{N}(\xi,0,s)\\ \mathbf{N}(\xi,z,s)\end{bmatrix}\mathbf{L}. \qquad \text{(S47)}$$

Because $\mathbf{\Lambda}$ and $\mathbf{V}$ can be expressed by the same constants $l_i$, combining S41 and S47 to eliminate $\mathbf{L}$ yields the relationship between the state vectors through the stiffness matrix $\mathbf{K}$ :

$$\begin{bmatrix}-\mathbf{V}(\xi,0,s)\\ \mathbf{V}(\xi,z,s)\end{bmatrix}=\mathbf{K}\begin{bmatrix}\mathbf{\Lambda}(\xi,0,s)\\ \mathbf{\Lambda}(\xi,z,s)\end{bmatrix}. \qquad \text{(S48)}$$

The elements in $\mathbf{K}$, which depends on the materials properties of the natural layer and the depth $z$ in the layer, also given in [3][a], are:

$$\mathbf{K}_{11}=\mathbf{K}_{44}=\frac{2G\xi b_2\left(4z\xi e_1^2 b_1-b_3 d_1 d_3\right)}{g}, \qquad \text{(S49)}$$

---

[a] A typo exists in the element $\mathbf{K}_{23}$ in reference [3] and is corrected here

$$\mathbf{K}_{12}=\mathbf{K}_{21}=-\mathbf{K}_{45}=-\mathbf{K}_{54}=\frac{2G\xi\left(4z^2\xi^2e_1^2b_1^2-Gb_3d_1^2\right)}{g},\qquad\text{(S50)}$$

$$\mathbf{K}_{13}=\mathbf{K}_{46}=-\frac{2G\beta a\xi\left(4z\xi e_1b_1\left(pe_2d_1-\xi e_1d_2\right)+b_3d_1\left(\xi d_2d_3-pd_1d_4\right)\right)}{sgd_2},\text{(S51)}$$

$$\mathbf{K}_{14}=\mathbf{K}_{41}=\frac{4G\xi e_1b_2\left(b_3d_1-z\xi b_1d_3\right)}{g},\qquad\text{(S52)}$$

$$\mathbf{K}_{15}=\mathbf{K}_{51}=-\mathbf{K}_{24}=-\mathbf{K}_{42}=\frac{-4Gz\xi^2e_1b_1b_2d_1}{g},\qquad\text{(S53)}$$

$$\mathbf{K}_{16}=\mathbf{K}_{43}=\frac{-4G\beta a\xi\left(z\xi e_1b_1\left(\xi d_2d_3-pd_1d_4\right)+b_3d_1\left(pe_2d_1-\xi e_1d_2\right)\right)}{sgd_2},\quad\text{(S54)}$$

$$\mathbf{K}_{22}=\mathbf{K}_{55}=\frac{-2G\xi b_2\left(4z\xi e_1^2b_1+b_3d_1d_3\right)}{g},\qquad\text{(S55)}$$

$$\mathbf{K}_{23}=-\mathbf{K}_{56}=\frac{2G\beta a\xi\left(4z\xi pe_1b_1f+4pe_1e_2b_3d_1+b_3d_1\left(\xi d_1d_2-pd_3d_4\right)\right)}{sgd_2},\text{(S56)}$$

$$\mathbf{K}_{25}=\mathbf{K}_{52}=\frac{4G\xi e_1b_2\left(b_3d_1+z\xi b_1d_3\right)}{g},\qquad\text{(S57)}$$

$$\mathbf{K}_{26}=-\mathbf{K}_{53}=\frac{4G\beta a\xi\left(4z\xi pe_1^2e_2b_1+z\xi e_1b_1\left(\xi d_1d_2-pd_3d_4\right)+pb_3d_1f\right)}{sgd_2},\text{(S58)}$$

$$\mathbf{K}_{33}=\mathbf{K}_{66}=\frac{-\kappa pd_4}{sd_2},\qquad\text{(S59)}$$

$$\mathbf{K}_{36}=\mathbf{K}_{63}=\frac{2\kappa pe_2}{sd_2},\qquad\text{(S60)}$$

$$\mathbf{K}_{13}=\mathbf{K}_{32}=\mathbf{K}_{34}=\mathbf{K}_{35}=\mathbf{K}_{61}=\mathbf{K}_{62}=\mathbf{K}_{64}=\mathbf{K}_{65}=0,\qquad\text{(S61)}$$

where $p=\left(s/a+\xi^2\right)^{1/2}$, $b_1=\lambda+G$, $b_2=\lambda+2G$, $b_3=\lambda+3G$, $e_1=e^{-z\xi}$, $e_2=e^{-zp}$, $d_1=1-e_1^2$, $d_2=1-e_2^2$, $d_3=1+e_1^2$, $d_4=1+e_2^2$, $f=e_2d_3-e_1d_4$, $g=4z^2\xi^2e_1^2b_1^2-b_3^2d_1^2$.

More generally, for a multi-layered medium, the relationship between planes at depth $h_i$ and $h_{i+1}$ within a finite layer made of a single material is:

$$\begin{bmatrix}-\mathbf{V}\left(\xi,h_i,s\right)\\ \mathbf{V}\left(\xi,h_{i+1},s\right)\end{bmatrix}=\mathbf{K}^{(k)}\begin{bmatrix}\mathbf{\Lambda}\left(\xi,h_i,s\right)\\ \mathbf{\Lambda}\left(\xi,h_{i+1},s\right)\end{bmatrix},\qquad\text{(S62)}$$

where $\mathbf{K}^{(k)}$ is the stiffness matrix of the $k$-th layer. Therefore, for our tissue phantom, depicted in Fig. S2, the relationship between the state vectors at depth $z_A = 0$, $z_C$, $z_D$, $z_E$ can be expressed as follows:

$$\begin{bmatrix} -\mathbf{V}\left(\xi, 0^+, s\right) \\ \mathbf{V}\left(\xi, z_c^-, s\right) \end{bmatrix} = \mathbf{K}^{(1,2)} \begin{bmatrix} \mathbf{\Lambda}\left(\xi, 0^+, s\right) \\ \mathbf{\Lambda}\left(\xi, z_c^-, s\right) \end{bmatrix}, \qquad \text{(S63)}$$

$$\begin{bmatrix} -\mathbf{V}\left(\xi, z_c^+, s\right) \\ \mathbf{V}\left(\xi, z_D^-, s\right) \end{bmatrix} = \mathbf{K}^{(3,4)} \begin{bmatrix} \mathbf{\Lambda}\left(\xi, z_c^+, s\right) \\ \mathbf{\Lambda}\left(\xi, z_D^-, s\right) \end{bmatrix}, \qquad \text{(S64)}$$

$$\begin{bmatrix} -\mathbf{V}\left(\xi, z_D^+, s\right) \\ \mathbf{V}\left(\xi, z_E^-, s\right) \end{bmatrix} = \mathbf{K}^{(5)} \begin{bmatrix} \mathbf{\Lambda}\left(\xi, z_D^+, s\right) \\ \mathbf{\Lambda}\left(\xi, z_E^-, s\right) \end{bmatrix}, \qquad \text{(S65)}$$

where $\mathbf{K}^{(1,2)}$, $\mathbf{K}^{(3,4)}$, $\mathbf{K}^{(5)}$ are the stiffness matrices corresponding to the combined layers 1 and 2, and 3 and 4, and the layer 5. Layers 1 and 2, and 3 and 4 can be combined as we assume the same materials properties for those PMDS layers. For the top and bottom half-spaces, the relationship between the state vectors can be simplified to:

$$\mathbf{V}\left(\xi, 0^-, s\right) = \mathbf{K}^{(-\infty)} \mathbf{\Lambda}\left(\xi, 0^-, s\right), \qquad \text{(S66)}$$

and:

$$-\mathbf{V}\left(\xi, z_E^+, s\right) = \mathbf{K}^{(+\infty)} \mathbf{\Lambda}\left(\xi, z_E^+, s\right), \quad \text{(S67)}$$

which consider no stress, displacement, heat transfer, or temperature rise at infinity:

$$\mathbf{V}\left(\xi, -\infty, s\right) = \mathbf{V}\left(\xi, +\infty, s\right) = \mathbf{\Lambda}\left(\xi, -\infty, s\right) = \mathbf{\Lambda}\left(\xi, +\infty, s\right) = 0. \qquad \text{(S68)}$$

$\mathbf{K}^{(+\infty)}$ and $\mathbf{K}^{(-\infty)}$ are $3 \times 3$ submatrices of $\mathbf{K}$ with:

$$\mathbf{K}^{(+\infty)} = \lim_{z \to +\infty} \begin{bmatrix} \mathbf{K}_{11} & \mathbf{K}_{12} & \mathbf{K}_{13} \\ \mathbf{K}_{21} & \mathbf{K}_{22} & \mathbf{K}_{23} \\ \mathbf{K}_{31} & \mathbf{K}_{32} & \mathbf{K}_{33} \end{bmatrix} = \begin{bmatrix} \mathbf{K}_{11}^{(+\infty)} & \mathbf{K}_{12}^{(+\infty)} & \mathbf{K}_{13}^{(+\infty)} \\ \mathbf{K}_{21}^{(+\infty)} & \mathbf{K}_{22}^{(+\infty)} & \mathbf{K}_{23}^{(+\infty)} \\ \mathbf{K}_{31}^{(+\infty)} & \mathbf{K}_{32}^{(+\infty)} & \mathbf{K}_{33}^{(+\infty)} \end{bmatrix}, \qquad \text{(S69)}$$

$$\mathbf{K}_{11}^{(+\infty)} = \mathbf{K}_{22}^{(+\infty)} = 2G\xi b_2 / b_3, \qquad \text{(S70)}$$

$$\mathbf{K}_{12}^{(+\infty)} = \mathbf{K}_{21}^{(+\infty)} = -2G^2 \xi / b_3, \qquad \text{(S71)}$$

$$\mathbf{K}_{13}^{(+\infty)} = \mathbf{K}_{23}^{(+\infty)} = 2G\beta a\xi\left(\xi - p\right) / sb_3, \qquad \text{(S72)}$$

$$\mathbf{K}_{31}^{(+\infty)} = \mathbf{K}_{32}^{(+\infty)} = 0, \qquad \text{(S73)}$$

$$\mathbf{K}_{33}^{(+\infty)} = -\kappa p / s, \quad \text{(S74)}$$

and:

$$\mathbf{K}^{(-\infty)} = \lim_{z\to+\infty}\begin{bmatrix}\mathbf{K}_{44} & \mathbf{K}_{45} & \mathbf{K}_{46}\\ \mathbf{K}_{54} & \mathbf{K}_{55} & \mathbf{K}_{56}\\ \mathbf{K}_{64} & \mathbf{K}_{65} & \mathbf{K}_{66}\end{bmatrix} = \begin{bmatrix}\mathbf{K}_{11}^{(+\infty)} & -\mathbf{K}_{12}^{(+\infty)} & \mathbf{K}_{13}^{(+\infty)}\\ -\mathbf{K}_{21}^{(+\infty)} & \mathbf{K}_{22}^{(+\infty)} & -\mathbf{K}_{23}^{(+\infty)}\\ \mathbf{K}_{31}^{(+\infty)} & -\mathbf{K}_{32}^{(+\infty)} & \mathbf{K}_{33}^{(+\infty)}\end{bmatrix}. \quad \text{(S75)}$$

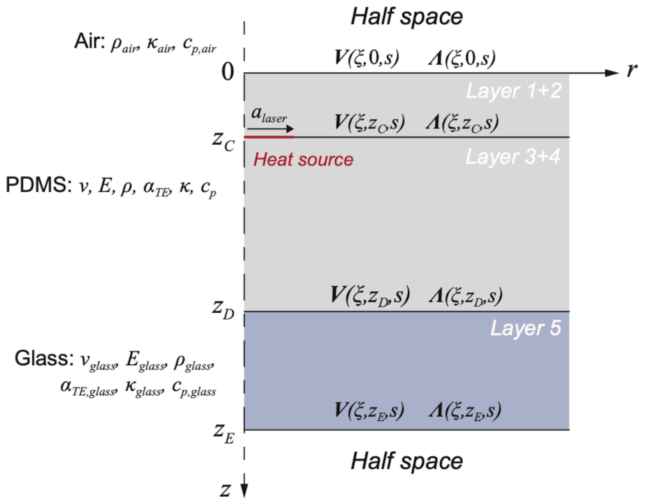


Fig. S2. Stress ( $\mathbf{V}$ ) and displacement ( $\mathbf{\Lambda}$ ) vectors in the Hankel-Laplace domain for the layered system. PDMS Layers 1 and 2, and 3 and 4 are combined. The surface heat source is located at depth $z_C$.

The continuity conditions at interfaces between layers, in the absence of external forces or heat sources, implies that:

$$\mathbf{V}\left(\xi,0^-,s\right) = \mathbf{V}\left(\xi,0^+,s\right), \quad \text{(S76)}$$

$$\mathbf{V}\left(\xi,z_D{}^-,s\right) = \mathbf{V}\left(\xi,z_D{}^+,s\right), \quad \text{(S77)}$$

$$\mathbf{V}\left(\xi,z_E{}^-,s\right) = \mathbf{V}\left(\xi,z_E{}^+,s\right), \quad \text{(S78)}$$

$$\mathbf{\Lambda}\left(\xi,0^-,s\right) = \mathbf{\Lambda}\left(\xi,0^+,s\right), \quad \text{(S79)}$$

$$\mathbf{\Lambda}\left(\xi,z_C{}^-,s\right) = \mathbf{\Lambda}\left(\xi,z_C{}^+,s\right), \quad \text{(S80)}$$

$$\mathbf{\Lambda}\left(\xi,z_D{}^-,s\right) = \mathbf{\Lambda}\left(\xi,z_D{}^+,s\right), \quad \text{(S81)}$$

$$\mathbf{\Lambda}\left(\xi,z_E{}^-,s\right) = \mathbf{\Lambda}\left(\xi,z_E{}^+,s\right). \quad \text{(S82)}$$

At the depth $z_c$, the boundary condition is:

$$\mathbf{V}\left(\xi,z_C{}^+,s\right) = \mathbf{V}\left(\xi,z_C{}^-,s\right) + \mathbf{F}\left(\xi,z_C,s\right), \quad \text{(S83)}$$

where the heat source is located. The external heat is source is expressed as:

$$\mathbf{F}(\xi, z_C, s) = \begin{bmatrix} 0 & 0 & \widetilde{Q_e}^{0}(\xi, z_C, s) \end{bmatrix}, \qquad \text{(S84)}$$

where $\widetilde{Q_e}^{0}(\xi, z_C, s)$ is the HL transform of the external heat flow $Q_e(r,t)$ brought by laser irradiation. Following the surface heat source description (equation 15 in the main text):

$$q_e(r,t) = q_0\,\phi_{laser}(r,t), \quad \text{(S85)}$$

where:

$$\phi_{laser}(r,t) = \left(H(t) - H(t-t_0)\right) \times \left( H(a_{laser} - r) * \frac{1}{2\pi\sigma_{defocus}^{2}} e^{-\frac{r^2}{2\sigma_{defocus}^{2}}} \right), \quad \text{(S86)}$$

we obtain:

$$\widetilde{Q_e}^{0}(\xi, z_C, s) = \frac{1}{s}\widetilde{q_e}^{0}(\xi, s) = \frac{q_0 a J_1(\xi a)\left(1 - e^{-t_0 s}\right)}{\xi s^2} e^{-\frac{\sigma_{defocus}^2 \xi^2}{2}}. \qquad \text{(S87)}$$

We apply additional boundary conditions, which includes the absence of stress at the air/glass and air/PDMS boundaries. We, however, consider heat transfer at the top interface and ignore it at the bottom interface, which is far from the measurement planes so:

$$\mathbf{V}(\xi, 0, s) = \left[0, 0, \tilde{Q}^{0}(\xi, 0, s)\right]^{T}, \quad \text{(S88)}$$

$$\mathbf{V}(\xi, z_E, s) = 0, \quad \text{(S89)}$$

leading to:

$$\mathbf{K}^{(+\infty)} = 0, \qquad \text{(S90)}$$

$$\mathbf{K}^{(-\infty)} = \begin{bmatrix} 0 & 0 & 0 \\ 0 & 0 & 0 \\ 0 & 0 & -\kappa p / s \end{bmatrix}. \text{(S91)}$$

Although the surface heat transfer with air is relatively small, it will be important to model it in the eye as the retina is in contact with the vitreous, which has a higher thermal conductivity than air. Rewriting $\mathbf{K}^{(1,2)}$ as $\mathbf{K}^{(1,2)} = \begin{bmatrix} \mathbf{A}^{(1,2)} & \mathbf{B}^{(1,2)} \\ \mathbf{C}^{(1,2)} & \mathbf{D}^{(1,2)} \end{bmatrix}$ and likewise for $\mathbf{K}^{(3,4)}$ and $\mathbf{K}^{(5)}$, we can derive the state vectors of interest $\mathbf{\Lambda}(\xi, 0, s)$ and $\mathbf{\Lambda}(\xi, z_c, s)$ from equations S63–66 and S76–83:

$$\mathbf{\Lambda}(\xi, z_c, s) = -\left( -\mathbf{C}^{(1,2)} \left( \left( \mathbf{K}^{(-\infty)} + \mathbf{A}^{(1,2)} \right)^{-1} \mathbf{B}^{(1,2)} \right) + \mathbf{D}^{(1,2)} + \mathbf{A}^{(3,4)} + \mathbf{B}^{(3,4)}\mathbf{R} \right)^{-1} \mathbf{F}, \qquad \text{(S92)}$$

with:

$$\mathbf{R} = -\left( \mathbf{D}^{(3,4)} + \mathbf{A}^{(5)} - \mathbf{B}^{(5)} \left( \left( \mathbf{D}^{(5)} \right)^{-1} \mathbf{C}^{(5)} \right) \right)^{-1} \mathbf{C}^{(3,4)}, \qquad \text{(S93)}$$

and:

$$\mathbf{\Lambda}(\xi,0,s) = -\left(\mathbf{K}^{(-\infty)} + \mathbf{A}^{(1,2)}\right)^{-1} \mathbf{B}^{(1,2)} \mathbf{\Lambda}(\xi, z_c, s). \quad \text{(S94)}$$

For an intermediate plane located at a depth $z_I$ between the planes A and C, the state vector is:

$$\mathbf{\Lambda}(\xi,z_I,s) = -\left(\mathbf{D}^{(1,2)}(\xi,z_I,s) + \mathbf{A}^{(1,2)}(\xi,z_c - z_I,s)\right)^{-1} \left(\mathbf{C}^{(1,2)}(\xi,z_I,s)\mathbf{\Lambda}(\xi,0,s) + \mathbf{B}^{(1,2)}(\xi,z_c - z_I,s)\mathbf{\Lambda}(\xi,z_c,s)\right) . \quad \text{(S95)}$$

We recall that to calculate $\overline{\mathrm{AA}}(r,t)$ we need to calculate $u_z(r,z_A,t)$ at plane A for which $\mathbf{\Lambda}(\xi,0,s)$ is used. To calculate $\overline{\mathrm{AC}}(r,t)$, we not only need $u_z(r,z_A,t)$ and $u_z(r,z_C,t)$, which require $\mathbf{\Lambda}(\xi,0,s)$ and $\mathbf{\Lambda}(\xi,z_c,s)$, but also $\overline{\Delta n}$ or the average temperature rise $\theta$ between the planes A and C, which requires $\mathbf{\Lambda}(\xi,z_I,s)$. We determined that the calculating this solution vector for 40 evenly-spaced depths between A and C was sufficient to ensure convergence of $\overline{\Delta n}$.

Having established the complete solutions for the displacements and the temperature rise in the HL domain, the spatial and time domains solutions can be found by numerical inversion of the Hankel and Laplace transforms, where both can be treated independently. For the Laplace transform, the Abate-Whitt framework [4] is a commonly used set of methods in which the inverse transform $f$ is described by a linear combination of transformed values:

$$f(t) \approx \sum_{k=1}^{N} \frac{\eta_k}{t} \bar{f}\left(\frac{\beta_k}{t}\right),\ t > 0, \quad \text{(S96)}$$

where $\bar{f}$ is the Laplace transform of $f$, the nodes $\beta_k$ and weights $\eta_k$ are $N$-dependent real or complex numbers. The precision of the inversion increases with $N$. Common methods in this framework include the Gaver-Stehfest, the Euler, and the Talbot method but all have been shown to be numerically unstable when inversing discontinuous functions, such as the square function [5,6]. We used instead a more recently developed method, which uses concentrated matrix exponential (CME) distribution and demonstrates remarkable stability and accuracy for discontinuous functions [6]. The $\beta_k$ and $\eta_k$ parameters are made available by the authors of [6].

The inverse Hankel transform of $\hat{f}(\xi)$ is defined by the integral:

$$f(r) = \int_0^{+\infty} \hat{f}(\xi) J_m(\xi r)\xi d\xi, \quad \text{(S97)}$$

which can be split into intervals [ $\xi_i, \xi_{i+1}$ ]:

$$\int_0^{+\infty} \hat{f}(\xi) J_m(\xi r)\xi d\xi = \sum_{i=0}^{\infty} \int_{\xi_i}^{\xi_{i+1}} \hat{f}(\xi) J_m(\xi r)\xi d\xi \approx \sum_{i=0}^{N_B} \int_{\xi_i}^{\xi_{i+1}} \hat{f}(\xi) J_m(\xi r)\xi d\xi, \quad \text{(S98)}$$

where $\xi_i$ represent the zero points of the Bessel function $J_m$. The integrals over the intervals [ $\xi_i, \xi_{i+1}$ ] can then be approximated using the Gauss-Legendre quadrature with:

$$\int_{\xi_i}^{\xi_{i+1}} \hat{f}(\xi) J_m(\xi r)\xi d\xi = \frac{\xi_{i+1} - \xi_i}{2} \sum_{p=1}^{N_{GL}} w_p \hat{f}(\beta_k) J_m(r\beta_k)\beta_k, \quad \text{(S99)}$$

$$\beta_k = \left(\frac{\xi_{i+1} - \xi_i}{2}\right) r_p + \left(\frac{\xi_{i+1} + \xi_i}{2}\right), \qquad \text{(S100)}$$

where $w_p$ and $r_p$ are the Gaussian weights and nodes, respectively [7]. The accuracy of the numerical integration increases with the number of Bessel intervals considered ( $N_B$ ) and the quadrature order $N_{GL}$ (see the following section for numerical inversion convergence study).

### 3. Stiffness matrix method (SMM) validation using FEM simulations

To validate the SMM calculations, we used an axisymmetric finite-element model of the tissue phantom implemented in the finite element method (FEM) software COMSOL. The model included all the phantom layers listed in Table 1 and a finite air layer at the top of the assembly, in contact with plane A. Contrary to SMM, the model must be of finite size in all dimensions, so the air layer was chosen to have a thickness 500-μm and the width of the domain was 2000 μm ($10\times$ the heating beam radius). The bottom interface (glass) was considered to be thermally isolated. The temperature was set at room temperature on the other boundaries.

The boundary heat source was modeled following the Beer-Lambert law as described in the Materials and Methods, equation 15:

$$q_e(r,t) = \frac{P_0}{\pi a^2} \phi_{laser}(r,t)\left(1 - e^{-\mu_a l}\right). \quad \text{(S101)}$$

We here omitted for validation purposes the Gaussian blur in the spatial profile of the beam and considered a square pulse in time with a top-hat profile in space.

$$\phi_{laser}(r,t) = \left(H(t) - H(t - t_0)\right) \times H\left(a_{laser} - r\right). \qquad \text{(S102)}$$

The validation was performed using the fitted material parameters listed in Table 2. The COMSOL model included the solid mechanics module with linear elasticity equations, the heat transfer module, and the thermal expansion multi-physics module. The mesh was defined using the extremely-fine physics-controlled mesh provided by COMSOL to ensure element size convergence. In Fig. S3a, we compare the vertical displacements at the center ($r$ = 0) of plane A ($z = z_A$) obtained from COMSOL simulations and calculated by SMM using high-order inverse integrations for both the Hankel and Laplace inverse transforms (20 Bessel intervals (B=20), 20-point Gauss-Legendre quadrature (G=20), 20$^{th}$ order inverse Laplace with concentrated matrix exponential distributions (C=20)). We thus verify the good match between the calculation methods. We note short-lived oscillations (μs timescale) in COMSOL results corresponding to the establishment of the mechanical equilibrium (wave reflections) that are absent in SMM results. The time-dependent term in the wave equation is ignored in the governing equations for SMM (equations S13 and S14) as the wave dynamics are too fast to be captured using the OCT system. We also note a faster recovery of $u_z$ in COMSOL, which is attributed to boundary effects in COMSOL (finite air thickness and finite width).

Having verified the accuracy of the SMM calculations at high integration orders, we studied the effect of integration orders of the precision on the results. Since the computation time depends on these orders, we aimed to select low orders giving reasonable accuracy and precision. The error between the high-order calculations (B=20, G=20, C=20) and lower-order calculations is shown in Fig. S3b. We estimated that convergence below 1% error occurs for B≥5, G≥6, and C≥7. These orders (B=5, G=6, C=7) were therefore selected for the fitting procedure.

For a simulated duration of 200 ms, with 0.1-ms time intervals, the computation time for COMSOL was about 30 minutes. Using SMM and selected orders, for the same time points (2001 points), and for one radial point ($r$ = 0), the computation took 70 seconds on a laptop computer, which demonstrates the increased computation efficiency using SMM.

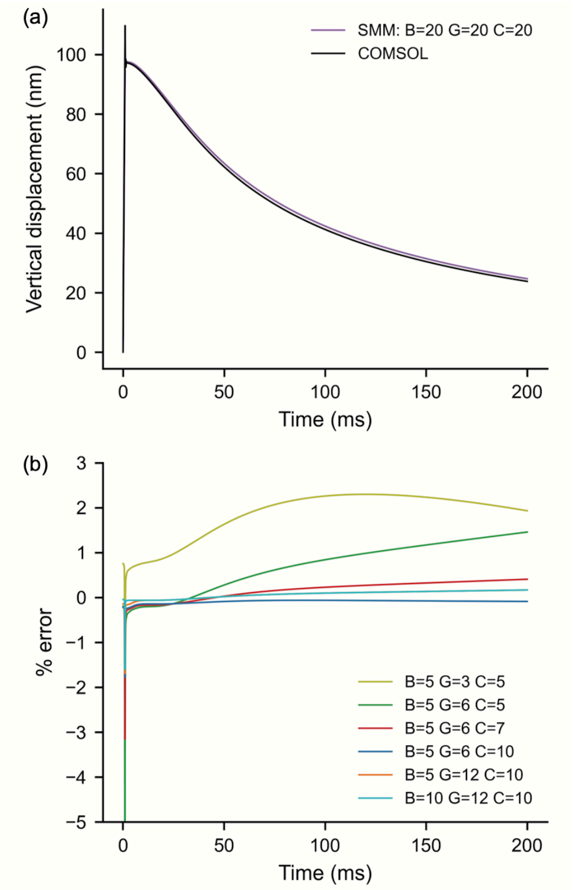


Fig S3. (a) Vertical displacements $u_z$ calculated at ($r$ = 0, $z_A$ = 0) with COMSOL and stiffness matrix method (SMM) with 20 Bessel intervals (B=20) and a 20-point Gauss-Legendre quadrature of 20 (G=20) for the inverse Hankel transform and a concentrated matrix exponential order of 20 (C=20) for the inverse Laplace transform. (b) Error between high-order inverse HL transform (B=20, G=20, C=20) and lower-order inverse transforms.

### 4. Elastic modulus effects on thermal expansion

We evaluated the influence of the elastic modulus on the thermo-mechanical response by running SMM calculations varying the elastic modulus by a factor of 2 from the initial of $E$ = 1.3 MPa found in the literature [8]. Results are shown in Fig. S4. As one can expect, the elastic modulus has no effect on the temperature rise at the absorption layer (Fig. S4a). Additionally, the effect is also negligible on the vertical displacements $u_z$, as illustrated in Fig. S4b, which shows the vertical displacement calculated at ($r$ = 0, $z$ = 0). We could therefore set the elastic modulus at 1.3 MPa and not fit it.

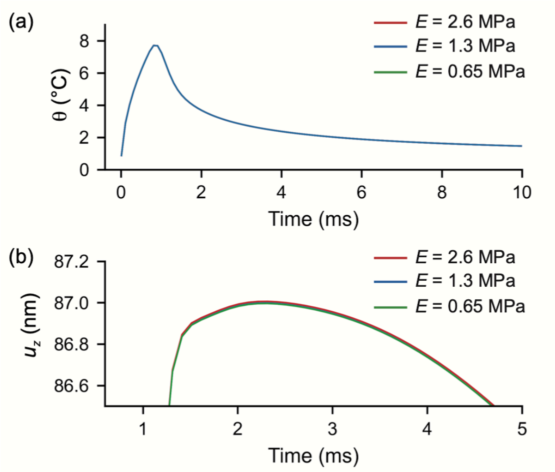


Fig. S4. Effects of the PDMS elastic modulus on the tissue model response. (a) $E$ has no effect on the temperature rise ($\theta$). All curves overlap and are undistinguishable. (b) $E$ has little effect on the vertical displacement response.

### 5. Material property fitting procedure

The raw OCT data after calculations of the ΔOPLs, which removes the $z$ dimension, has 521 points in radius and 5001 points in time. This corresponds to a radial extent of ±1150 um and a duration of 500 ms. As we fitted the $\overline{\mathrm{AA}}$ data first, we cropped the dataset radially to a smaller window of ± 500 μm, which encompassed the highest SNR data points, which are most useful for fitting. We then performed a spatial average between positive and negative radii and a temporal rolling average using a rolling window of 5 time points. Because the rolling average correlate noise in the rolling window, we extracted a subset of time points to preserve uncorrelated-noise data. We also cropped the data in time starting shortly after the end of the heating pulse (1.5 ms), after which the data varied slowly so that the rolling average did not erase high-frequency features. At this delay, the surface and the volumetric heat sources can be considered equivalent. The $\overline{\mathrm{AA}}$ dataset at this stage was composed of 111 points in space and 961 points in time. As explained in the main text, by fitting $\overline{\mathrm{AA}}$ we can determine the parameters $\kappa$, $\sigma_{defocus}$, $a_{laser}$ and the proportionality factor $\gamma$ to the product $\alpha_{TE} \times q_0$ with $\left(\alpha_{TE}\, q_0\right)_{fitted} = \gamma\left(\alpha_{TE}\, q_0\right)_{initial}$.

Because the fitting computation time depends on the number of experimental data points considered, we then aimed to determine the minimum number of points that would ensure accurate fitting. To do so, we down-selected an array of $n$-by-$n$ points (starting at $4 \times 4$ points), performed non-linear least square fitting on this dataset, got the fitted parameters, used the fitted parameters to calculate the modeled $\overline{\mathrm{AA}}$ on the full dataset domain ($111 \times 961$ points), and computed the root-mean-square error (RMSE) between the modeled and the full experimental $\overline{\mathrm{AA}}$ dataset. Then a new $n$-by-$n$ points dataset was extracted from the full dataset with a higher $n$ and fitting operations were repeated. It is expected that the RMSE will tend to decrease as we included more experimental data points (as $n$ increased) in the fitting. When the RMSE plateaued with increasing $n$, we could consider that additional data points would not bring added accuracy or precision. We therefore stopped at this $n$ value. We found that $22 \times 22$ $\overline{\mathrm{AA}}$ points were sufficient to obtain acceptable fitting.

To estimate the uncertainties on the parameters $\kappa$, $\sigma_{defocus}$, $a_{laser}$, and $\gamma$, the experimental data were bootstrapped to simulate 200 experimental datasets. The fitting was then repeated 200 times and we obtained 200-element arrays of fitted $\kappa$, $\sigma_{defocus}$, $a_{laser}$, and $\gamma$ values. The best fit values were calculated by averaging the 200 fitted values and the uncertainties were obtained from the standard deviations. The fitted values also served to evaluate the correlation between the parameters (see the following Parameter correlation section).

Using $\kappa$, $\sigma_{defocus}$, $a_{laser}$, and $\gamma$ found by fitting $\overline{\mathrm{AA}}$ we calculated the modeled $\overline{\mathrm{AC}}$ by applying the $\gamma$ factor to $q_0$. Here we aimed to find the independent and respective $\gamma_i$ values for $q_0$ and $\alpha_{TE}$ with:

$$\left(\alpha_{TE}\right)_{fitted} = \gamma_1 \left(\alpha_{TE}\right)_{initial} , \quad \text{(S103)}$$

$$\left(q_0\right)_{fitted} = \gamma_2 \left(q_0\right)_{initial} , \quad \text{(S104)}$$

$$\gamma = \gamma_1 \times \gamma_2 . \quad \text{(S105)}$$

Because $\overline{\mathrm{AC}}$ scales linearly with $q_0$ but not $\alpha_{TE}$, fitting a scaling factor between the modeled and the experimental $\overline{\mathrm{AC}}$ meant find the $\eta$ factor with:

$$\left(q_0\right)_{fitted} = \eta \gamma \left(q_0\right)_{initial} , \quad \text{(S106)}$$

which therefore allowed the determination of $\gamma_1$ and $\gamma_2$:

$$\gamma_2 = \eta\gamma , \quad \text{(S107)}$$

$$\gamma_1 = 1/\eta . \quad \text{(S108)}$$

$\overline{\mathrm{AC}}$ was also reduced to the same $22 \times 22$ points and bootstrapped 200 times to find the constant $\eta$ and its corresponding uncertainty. At this point the five fitting parameters $\kappa$, $\sigma_{defocus}$, $a_{laser}$, $q_0$ (therefore $\mu_a$) and $\alpha_{TE}$ were fully determined along with their uncertainties. The values are reported in Table 3. The full procedure is shown schematically in Fig. S5. The residuals obtained after fitting of $\overline{\mathrm{AA}}$ and $\overline{\mathrm{AC}}$ are shown in Fig. S6.

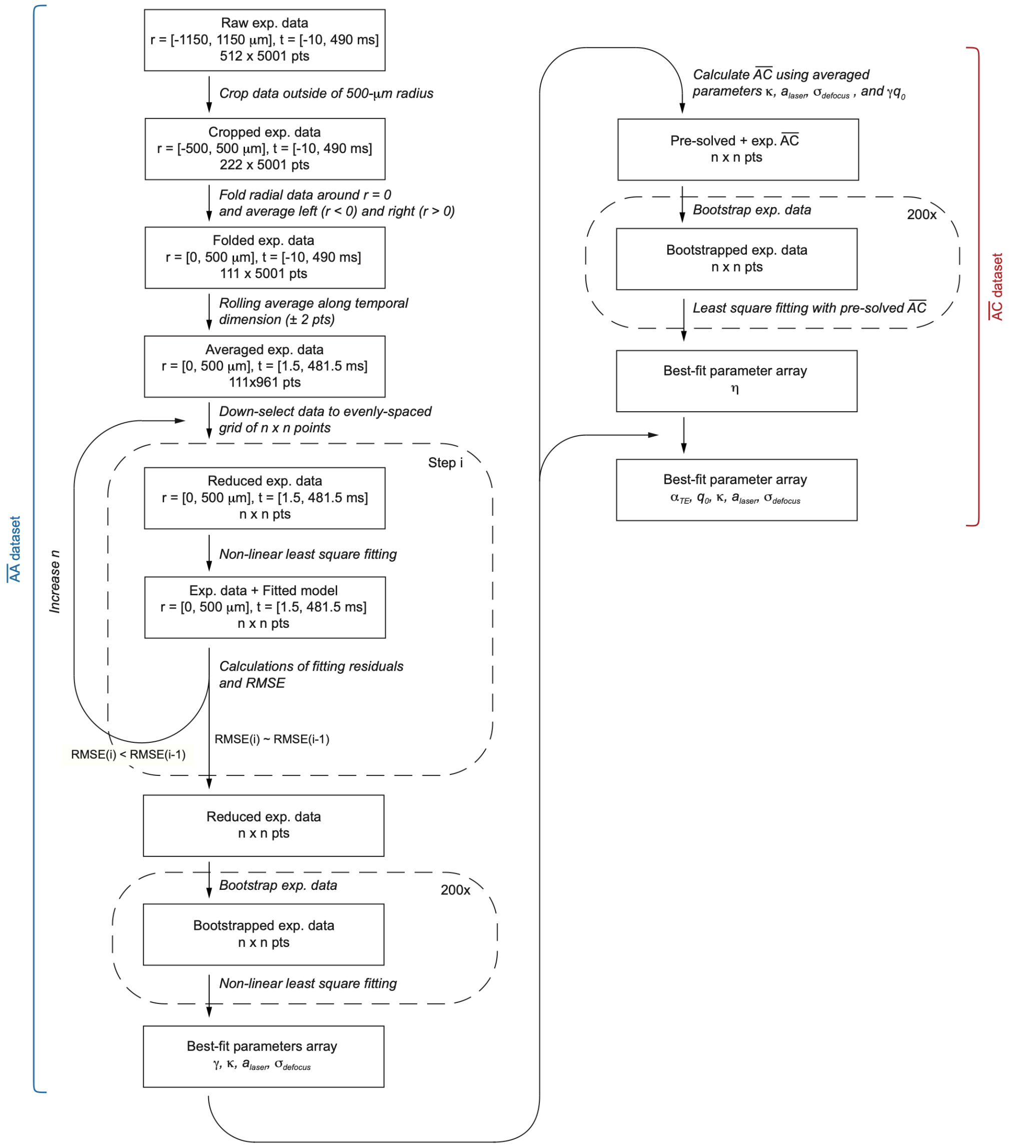


Fig. S5. Data reduction and fitting procedure.

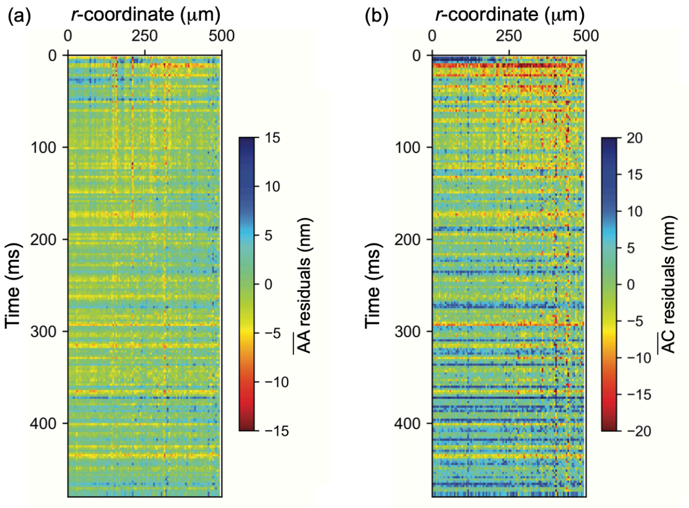


Fig. S6. Residuals between modeled and experimental ΔOPLs.

### 6. Parameter correlations

Through fitting of bootstrapped experimental data (200 times), it is possible to evaluate the cross-correlation between the fitting parameters. The correlation between parameters can be visualized by plotting the result of the 200 fits for all pairs of parameters (6 pairs for 4 variables) (Fig. S7). Low correlation is indicated by the absence of high-eccentricity ellipses not aligned with the parameter axes.

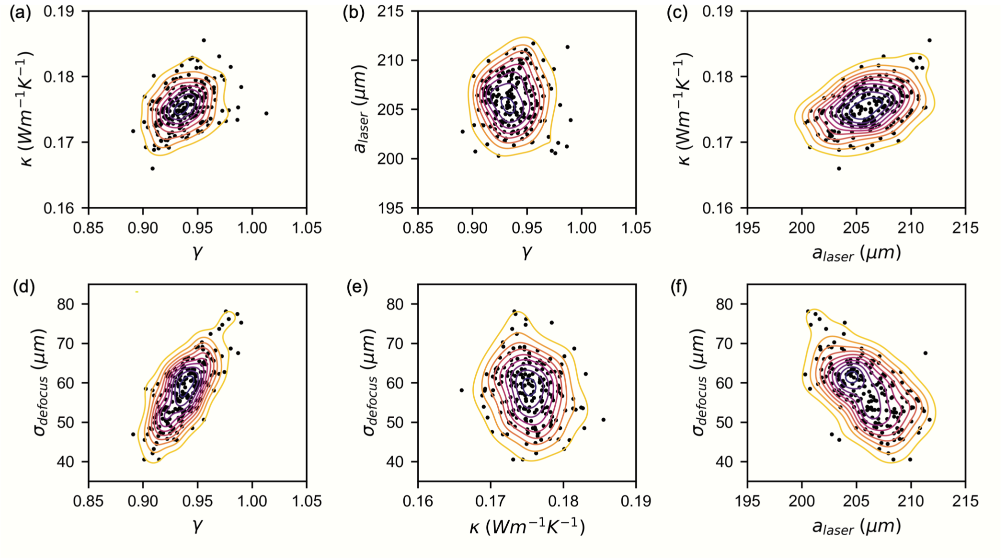


Fig. S7. Correlations between fitted parameter values obtained by bootstrapping. Local densities, indicated by the contour maps, were estimated using a Gaussian kernel.

### 7. Non-damaging treatment range

As described in the Materials and Methods section, the Arrhenius integral can be used to define the therapeutic window and determine the laser power titration. It can be expressed as [9]:

$$\Omega(r,t) = A\int_0^{\tau} e^{-\frac{E^*}{R\times T(t)}} dt, \quad \text{(S109)}$$

where $A$ ($1.6\times10^{55}$ $s^{-1}$) is the rate constant, $E^*$ (340 kJ/mol) is the activation energy, $R$ (8.314 $J.K^{-1}.mol^{-1}$) is the gas constant and $T$ is the temperature as a function of time. We here looked at the influence of the absorption coefficient on the integral value in order to estimate the precision required on this material parameter to allow non-damaging therapy. We ran COMSOL simulations similar to what is done to validate the stiffness matrix method with the tissue phantom (see Stiffness matrix method (SMM) validation using FEM simulations section in the SI). The retina was modeled as rat retina composed of multi-layered isotropic media whose top surface is in contact with a liquid (modeled as water) representing the vitreous fluid. The layer thicknesses and material properties used in the simulations are listed in Table S1 with the corresponding references. In COMSOL, the retinal layers were modeled as linear elastic layers and the vitreous a Newtonian fluid. Heat transfer was allowed at the liquid/solid interface. The external heat source was modeled following the Beer-Lambert law with the heat flux given by:

$$q_e(r,z,t) = \frac{P_0}{\pi a_{laser}^2}\phi_{laser}(r,t)\mu_a(z)e^{-\mu_a(z)z}, \quad \text{(S110)}$$

where $P_0$ is the incident laser power at the top of the absorbing layer, $a_{laser}$ is the radius of the top-hat intensity profile, $\phi_{laser}$ is the spatio-temporal profile of the laser beam described below and $\mu_a$ is the optical absorption coefficient. For the non-pigmented choroid (NPC) and the pigmented choroid (PC) layers, $P_0$ was calculated taking into account the absorption of the preceding layers. The beam spatio-temporal profile $\phi_{laser}$ was modeled as a square pulse in time with duration $t_0 = 10ms$ and top-hat profile in space with radius $a_{laser} = 200\mu m$, which are typical parameters used in retinal therapy [10]:

$$\phi_{laser}(r,t) = \left(H(t) - H(t-t_0)\right)\times H(a_{laser} - r), \quad \text{(S111)}$$

where $H$ is the Heaviside function. The initial absorption coefficient values $\mu_{a,\,0}$ are taken from literature [11]. Using initial values, $P_0$ was adjusted to 55 mW so that $\Omega \approx 0.5$ (about the center of the therapeutic window), which was calculated using the temperature course $T(t)$ at the hottest point in space (top of the retinal pigment epithelium (RPE) layer in depth, center of the beam in radius) (Fig S8a). The absorption coefficients $\mu_a$ was then varied and the Arrhenius integral was calculated (Fig S8b). Such study allows us to determine the acceptable precision on $\mu_a$ to remain in the therapeutic window. We found that a variation of about 20% from the initial values reached the limits of the window, which corresponded to a peak temperature difference of ±4°C.

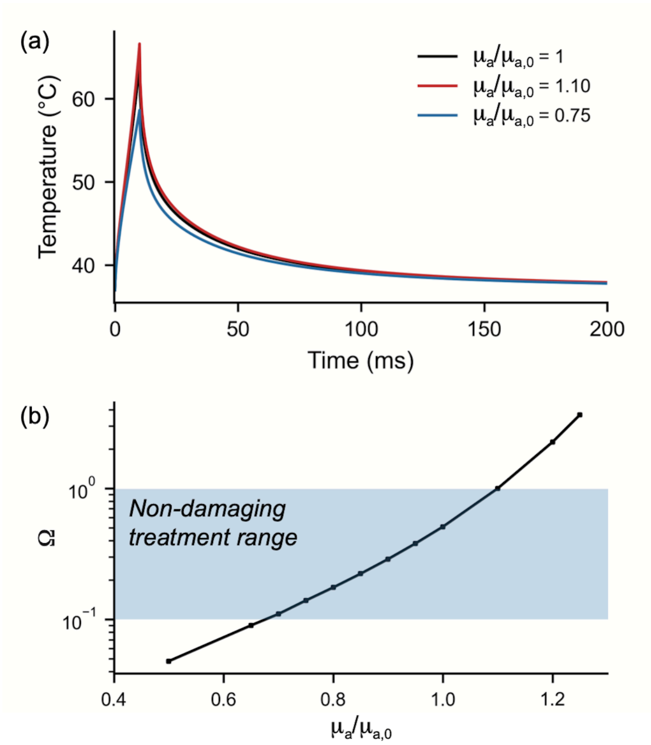


Fig. S8. (a) Temperature profiles at the top of the RPE layer and center of the beam for three absorption coefficient values. (b) Arrhenius integral calculated as a function of the absorption coefficient value $\mu_a$ relative to its initial value $\mu_{a,0}$. The non-damaging treatment range is $0.1 < \Omega < 1$.

Table S1. Retina model parameters used in COMSOL simulations for temperature calculations. NFL: nerve fiber layer, GCL: ganglion cell layer, IPL: inner plexiform layer, INL: inner nuclear layer, OPL: outer plexiform layer, ONL: outer nuclear layer, PR: photoreceptor layer, RPE: retinal, PC: pigmented choroid, NPC: non-pigmented choroid. *h*: layer thickness, *ρ:* density*, E:* Young's modulus, $\alpha_{TE}$*:* coefficient of thermal expansion, *ν:* Poisson's ratio, *κ*: coefficient of thermal conductivity, $c_p$: specific heat capacity, $\mu_a$: coefficient of optical absorption.

| | *h* (μm) | *ρ* (kg.m$^{-3}$) | *E* (kPa) | $\alpha_{TE}$ (K$^{-1}$) | *ν* | *κ* (W.m$^{-1}$.K$^{-1}$) | $c_p$ (J.kg$^{-1}$.K$^{-1}$) | $\mu_a$ (cm$^{-1}$)[b] |
|---|---|---|---|---|---|---|---|---|
| Water (vitreous) | 300 | 993[a] | NA | $3.45 \times 10^{-4}$ [a] | NA | 0.63[a] | 4180[a] | - |
| NFL-GCL | 30 [12] | | 1.3 [13] | | 0.47 [14] | 0.45 [15] | 3600 [16] | - |
| IPL | 30 [12] | | 1.3 [13] | | | | | - |
| INL | 20 [12] | | 2.7 [13] | | | | | - |
| OPL | 10 [12] | | 8 [13] | | | | | - |
| ONL | 65 [12] | | 2.7 [13] | | | | | - |
| PR | 40 [12] | | 26 [13] | | | | | - |
| RPE melanosomes | 1 [11] | | 5 [17] | | | | | $9.976 \times 10^3$ [11] |
| RPE basal | 3 [11] | | 5 [17] | | | | | - |
| PC | 20 [11] | | 30 [17] | | | | | $2.494 \times 10^3$ [11] |
| NPC | 30 [11] | | 30 [17] | | | | | 475 [11] |
| Sclera | 500 [11] | | 30 [17] | | | | 3200 [16] | - |

[a]www.engineeringtoolbox.com. Data taken at 37°C. [b]Small absorption coefficients (<1 cm$^{-1}$) are ignored. The references are indicated next to the values.
